# Whole genome sequence analysis of blood lipid levels in >66,000 individuals

**DOI:** 10.1101/2021.10.11.463514

**Authors:** Margaret Sunitha Selvaraj, Xihao Li, Zilin Li, Akhil Pampana, David Y Zhang, Joseph Park, Stella Aslibekyan, Joshua C Bis, Jennifer A Brody, Brian E Cade, Lee-Ming Chuang, Ren-Hua Chung, Joanne E Curran, Lisa de las Fuentes, Paul S de Vries, Ravindranath Duggirala, Barry I Freedman, Mariaelisa Graff, Xiuqing Guo, Nancy Heard-Costa, Bertha Hidalgo, Chii-Min Hwu, Marguerite R Irvin, Tanika N Kelly, Brian G Kral, Leslie Lange, Xiaohui Li, Martin Lisa, Steven A Lubitz, Ani W Manichaikul, Preuss Michael, May E Montasser, Alanna C Morrison, Take Naseri, Jeffrey R O’Connell, Nicholette D Palmer, Patricia A Peyser, Muagututia S Reupena, Jennifer A Smith, Xiao Sun, Kent D Taylor, Russell P Tracy, Michael Y Tsai, Zhe Wang, Yuxuan Wang, Bao Wei, John T Wilkins, Lisa R Yanek, Wei Zhao, Donna K Arnett, John Blangero, Eric Boerwinkle, Donald W Bowden, Yii-Der Ida Chen, Adolfo Correa, L Adrienne Cupples, Susan K Dutcher, Patrick T Ellinor, Myriam Fornage, Stacey Gabriel, Soren Germer, Richard Gibbs, Jiang He, Robert C Kaplan, Sharon LR Kardia, Ryan Kim, Charles Kooperberg, Ruth J. F. Loos, Karine Martinez, Rasika A Mathias, Stephen T McGarvey, Braxton D Mitchell, Deborah Nickerson, Kari E North, Bruce M Psaty, Susan Redline, Alexander P Reiner, Ramachandran S Vasan, Stephen S Rich, Cristen Willer, Jerome I Rotter, Daniel J Rader, Xihong Lin, NHLBI Trans-Omics for Precision Medicine (TOPMed) Consortium, Gina M Peloso, Pradeep Natarajan

**Affiliations:** Cardiovascular Research Center, Massachusetts General Hospital, Boston, MA, USA, 02114; Program in Medical and Population Genetics, Broad Institute of Harvard and MIT, Cambridge, MA, USA, 02142; Department of Medicine, Harvard Medical School, Boston, MA, USA, 02115; Department of Biostatistics, Harvard T.H. Chan School of Public Health, Boston, MA, USA, 02115; Department of Genetics, Perelman School of Medicine, University of Pennsylvania, Philadelphia, PA, USA, 19104; Department of Medicine, Perelman School of Medicine, University of Pennsylvania, Philadelphia, PA, USA,19104; Department of Epidemiology, University of Alabama at Birmingham School of Public Health; Cardiovascular Health Research Unit, Department of Medicine, University of Washington, Seattle, WA; Department of Medicine, Brigham and Women’s Hospital, Harvard Medical School; Department of Internal Medicine, National Taiwan University Hospital, Taipei, Taiwan; Institute of Population Health Sciences, National Health Research Institutes, Zhunan, 350, Taiwan; Department of Human Genetics and South Texas Diabetes and Obesity Institute, University of Texas Rio Grande Valley School of Medicine, Brownsville, TX 78520; Department of Medicine, Cardiovascular Division, Washington University School of Medicine, St. Louis, MO; Division of Biostatistics, Washington University School of Medicine, St. Louis, MO; Human Genetics Center, Department of Epidemiology, Human Genetics, and Environmental Sciences, School of Public Health, The University of Texas Health Science Center at Houston, Houston, Texas, USA; Department of Internal Medicine, Section on Nephrology, Wake Forest School of Medicine, Winston-Salem, NC, USA, 27157; Department of Epidemiology, UNC Chapel Hill; The Institute for Translational Genomics and Population Sciences, Department of Pediatrics, The Lundquist Institute for Biomedical Innovation at Harbor-UCLA Medical Center, Torrance, CA USA; Department of Neurology, Boston university School of Medicine; Section of Endocrinology and Metabolism, Department of Medicine, Taipei Veterans General Hospital; Department of Epidemiology, Tulane University School of Public Health and Tropical Medicine, New Orleans, Louisiana, US, 70112; Tulane University Translational Science Institute, New Orleans, Louisiana, US, 70112; Department of Medicine, Johns Hopkins University School of Medicine, Baltimore, MD, USA 21206; Division of Biomedical Informatics and Personalized Medicine, Department of Medicine; Department of Medicine, George Washington University, Washingron DC; Cardiovascular Disease Initiative, The Broad Institute of MIT and Harvard, Cambridge, MA 02124; Department of Public Health Sciences, Center for Public Health Genomics, University of Virginia, Charlottesville, VA USA; The Charles Bronfman Institute for Personalized Medicine, Icahn School of Medicine at Mount Sinai, New York, NY; Department of Medicine, University of Maryland School of Medicine, Baltimore, MD; Ministry of Health, Government of Samoa, Samoa; Department of Biochemistry, Wake Forest School of Medicine, Winston-Salem, NC, USA, 27157; Department of Epidemiology, University of Michigan, Ann Arbor, MI, USA 48109; Lutia i Puava ae Mapu i Fagalele, Apia, Samoa; Departments of Pathology & Laboratory Medicine and Biochemistry, Larner College of Medicine at the University of Vermont, Colchester, VT USA; Department of Laboratory Medicine and Pathology, University of Minneosta, Minneapolis, MN USA; Department of Biostatistics, Boston University School of Public Health, Boston, MA, USA, 02118; Department of Epidemiology, Universiyy of Iowa; Department of Medicine (Cardiology) and Department of Preventive Medicine, Northwestern University Feinberg School of Medicine, Chicago, IL; Dean’s Office, University of Kentucky College of Public Health; Lundquist Institute for Biomedical Innovation at Harbor-UCLA Medical Center; Department of Population Health Science, University of Mississippi Medical Center; McDonnell Genome Institute; Brown Foundation Institute of Molecular Medicine, McGovern Medical School, The University of Texas Health Science Center at Houston, Houston, Texas, 77225; Broad Institute, Cambridge, Massachusetts, 02142; New York Genome Center, New York, New York, 10013; Baylor College of Medicine Human Genome Sequencing Center, Houston, Texas, 77030; Department of Epidemiology and Population Health, Albert Einstein College of Medicine, Bronx NY 10461 USA; Division of Public Health Sciences, Fred Hutchinson Cancer Research Center, Seattle WA 98109; Psomagen; NNF Center for Basic Metabolic Research, University of Copenhagen, Cophenhagen, Denmark; Illumina; Department of Epidemiology, International Health Institute, Brown University, Providence RI; Geriatrics Research and Education Clinical Center, Baltimore Veterans Administration Medical Center, Baltimore, MD; University of Washington, Department of Genome Sciences, Seattle, Washington, 98195; Department of Epidemiology, University of Washington, Seattle, WA; Department of Health Services, University of Washington, Seattle, WA; Sections of Preventive medicine and Epidemiology, Cardiovascular medicine, Department of Medicine, Boston University School of Medicine; Department of Epidemiology, Boston University School of Public Health; Framingham Heart Study; University of Michigan, Internal Medicine, Ann Arbor, Michigan, 48109; Institute for Translational Medicine and Therapeutics, Perelman School of Medicine, University of Pennsylvania, Philadelphia, PA, USA, 19104; Department of Statistics, Harvard University, Cambridge, MA, USA, 02138

## Abstract

Plasma lipids are heritable modifiable causal factors for coronary artery disease, the leading cause of death globally. Despite the well-described monogenic and polygenic bases of dyslipidemia, limitations remain in discovery of lipid-associated alleles using whole genome sequencing, partly due to limited sample sizes, ancestral diversity, and interpretation of potential clinical significance. Increasingly larger whole genome sequence datasets with plasma lipids coupled with methodologic advances enable us to more fully catalog the allelic spectrum for lipids. Here, among 66,329 ancestrally diverse (56% non-European ancestry) participants, we associate 428M variants from deep-coverage whole genome sequences with plasma lipids. Approximately 400M of these variants were not studied in prior lipids genetic analyses. We find multiple lipid-related genes strongly associated with plasma lipids through analysis of common and rare coding variants. We additionally discover several significantly associated rare non-coding variants largely at Mendelian lipid genes. Notably, we detect rare *LDLR* intronic variants associated with markedly increased LDL-C, similar to rare *LDLR* exonic variants. In conclusion, we conducted a systematic whole genome scan for plasma lipids expanding the alleles linked to lipids for multiple ancestries and characterize a clinically-relevant rare non-coding variant model for lipids.

## Introduction

Discovery of rare alleles linked to plasma lipids (i.e., low-density lipoprotein cholesterol [LDL-C], high-density lipoprotein cholesterol [HDL-C], total cholesterol [TC], and triglycerides [TG]) continue to yield important translational insights toward coronary artery disease (CAD), including PCSK9 and ANGPTL3 inhibitors now available in clinical practice^1,2, 3, 4, 5^. The monogenic and polygenic bases of plasma lipids are well-suited to population-based discovery analyses and confer broader insights for genetic analyses of complex traits. We now evaluate numerous newly catalogued, largely rare, alleles never previously systematically analyzed with lipids.

Analyses of imputed array-derived genome-wide genotypes and whole exome sequences in hundreds of thousands of increasingly diverse individuals continue to uncover low-frequency protein-coding variants linked to lipids. Due to purifying selection, causal variants conferring large effects tend to occur relatively more recently, and are thus rare and often specific to families or communities^6^. Most discovery analyses for large-effect rare alleles have focused on the analysis of disruptive protein-coding variants given (1) well-recognized constraint in coding regions, (2) incomplete genotyping of rare non-coding sequence given relative sparsity of deep-coverage (i.e., >30X) whole genome sequencing (WGS), and (3) better prediction of coding versus non-coding sequence variation consequence^1, 7, 8, 9,10, 11, 12^. We recently described a statistical framework incorporating multi-dimensional reference datasets paired with genomic data to improve rare coding and non-coding variant analyses for WGS analysis of lipids and other complex traits^13, 14^. Furthermore, including individuals of non-European ancestry facilitates the discovery of both novel alleles at established loci as well as novel loci^14, 15, 16^.

Here, we examine the full allelic spectrum with plasma lipids using whole genome sequences and harmonized lipids from the National Heart, Lung, and Blood Institute (NHLBI) Trans-Omics for Precision Medicine (TOPMed) program^17, 18^. We studied 66,329 participants and 428 million variants across multiple ancestry groups – 44.48% European, 25.60% Black, 21.02% Hispanic, 7.11% Asian and 1.78% Samoan. We identified robust allelic heterogeneity at known loci with several novel variants at these loci; we additionally identified novel loci and pursued replication in independent cohorts (31.50% non-European samples). We then explored the association of genome-wide rare variants with lipids, with detailed explorations of rare coding and non-coding variant models at known Mendelian dyslipidemia genes. Our systemic effort yields new insights for plasma lipids provides a framework for population based WGS analysis of complex traits.

## Results

### Overview

We studied the TOPMed Freeze8 dataset of 66,329 samples from 21 studies and performed genome-wide association studies (GWAS) separately for the four plasma lipid phenotypes (i.e., LDL-C, HDL-C, TC and TG) using 28M individual autosomal variants (minor allele count [MAC] > 20) and aggregated rare autosomal variant (minor allele frequency [MAF] < 1%) association testing for 417M variants (**Fig. 1, Supplementary Fig. 1**). Secondarily, we associated individual variants with minor allele frequencies (MAF) > 0.01% within each ancestry group to detect ancestry-specific lipid-associated alleles. We intersected our results with currently published array-based GWAS results^15^ to identify novel associations with lipids. We performed replication analyses for the putative novel associations identified, in up to approximately 45,000 independent samples with array-based genotyping imputed to TOPMed. Finally, we conducted rare variant association studies as multiple aggregate tests across the genome to identify gene-specific functional categories and non-coding genomic regions influencing plasma lipid concentrations.

**Fig 1:**
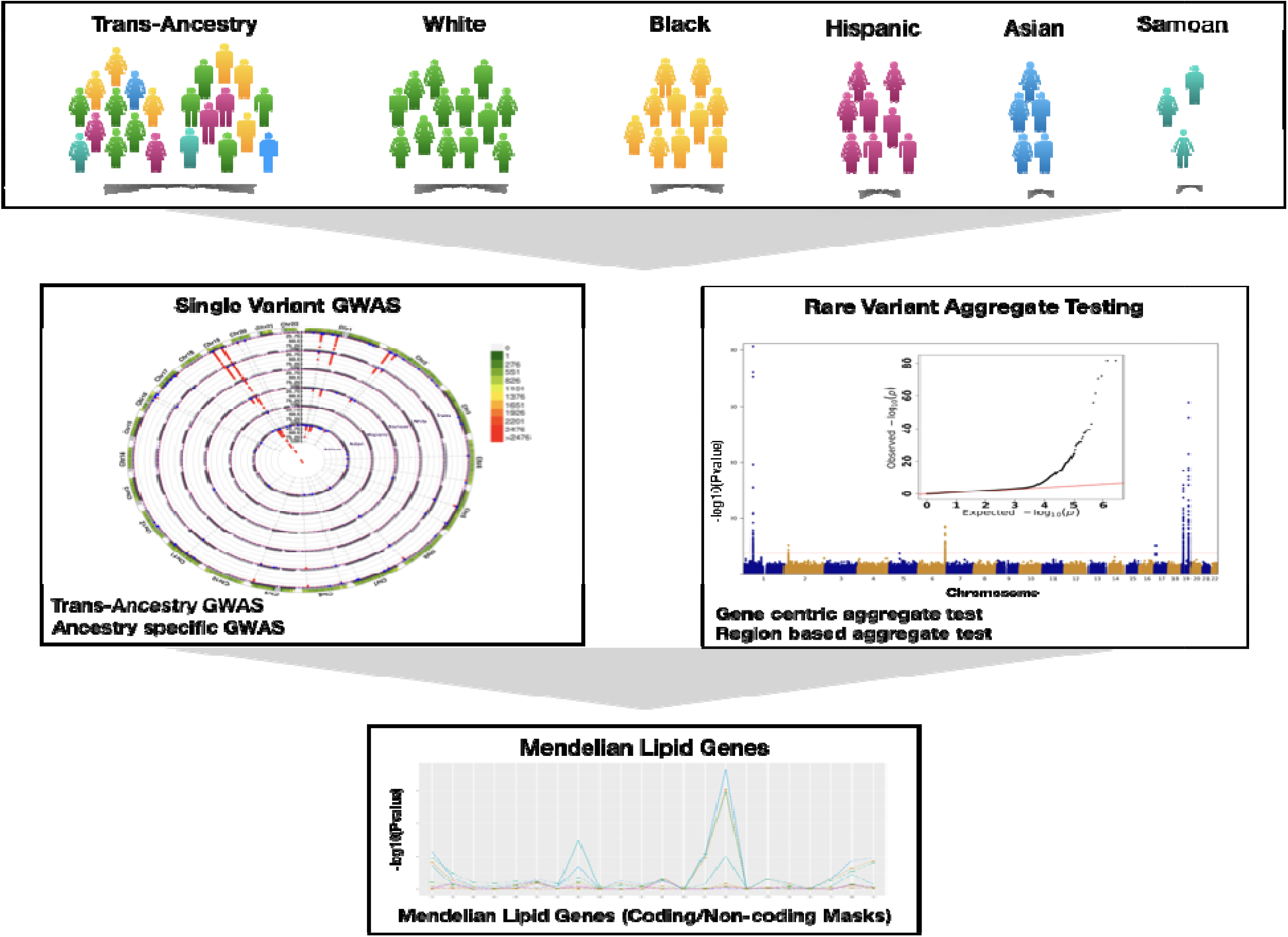
Overall study schematic. The analyses were conducted using the multi-ancestral TOPMed freeze8 data to associate whole genome sequence variation with lipid phenotypes (i.e., LDL-C, HDL-C, TC and TG). A total of 66,329 samples with lipids quantified data from five ancestry groups were analyzed. Single variant GWAS were carried out using SAIGE on the Encore platform using SNPs with MAC >20. Both trans-ancestry and ancestry-specific GWAS were conducted. Genome-wide rare variant (MAF < 1%) gene-centric and region-based aggregate tests were grouped and analyzed using STAARtopmed. Finally, single variant and rare variant associations at Mendelian dyslipidemia genes were investigated in further detail. TOPMed – Trans-Omics for Precision Medicine; HDL-C – High-Density Lipoprotein Cholesterol; LDL-C – Low-Density Lipoprotein Cholesterol; TC – Total Cholesterol; TG – Triglycerides; GWAS – Genome Wide Association Study; SAIGE – Scalable and Accurate Implementation of GEneralized mixed model; MAC – Minor Allele Count; MAF – Minor Allele Frequency; SNPs – Single nucleotide polymorphisms.

### TOPMed baseline characteristics

The TOPMed Informatics Research Center (IRC) and TOPMed Data Coordinating Center (DCC) performed quality control, variant calling, and calculated the relatedness of population structures of Freeze 8 data^17^. We studied 66,329 samples across 21 cohorts and 41,182 (62%) were female. The ancestry distribution was 29,502 (44.46%) White, 16,983 (25.60%) Black, 13,943 (21.02%) Hispanic, 4,719 (7.11%) Asian, and 1,182 (1.78%) Samoan (**Supplementary Table 1**). The mean (standard deviation [SD]) age of the full cohort was 53 (15.00) years which varied by cohort from 25 (3.56) years for Coronary Artery Risk Development in Young Adults (CARDIA) to 73 (5.38) years for Cardiovascular Health Study (CHS). The Amish cohort had a higher-than-average concentration of LDL-C (140 [SD 43] mg/dL) and HDL-C (56 [SD 16] mg/dL) as well as lower TG (median 63 [IQR 50] mg/dL) consistent with the known founder mutations in *APOB* and *APOC3*^7, 8, 14^. In the Women’s Health Initiative (WHI) cohort, the TC (230 [SD 41] mg/dL) and TG (median 129 [IQR 87] mg/dL) concentrations were higher than for other cohorts as previously described^12^. We accounted for lipid-lowering medications and fasting status and inverse rank normalized the phenotypes as before^12, 14^ which are further detailed in the **Methods**. The adjusted normalized lipid concentrations for the four lipids were similar across the cohorts.

A total of 428M variants passed the quality criteria with an average depth >30X in 22 autosomes. 202M variants were singletons, 417M were rare variants (MAF<1%), and 11M were common or low frequency variants (MAF>1%) with differences by cohort (**Supplementary Table 2**).

### Individual variant associations with lipids

Approximately 28M variants with MAC > 20 were individually associated with LDL-C, HDL-C, TC and TG. We used p-value < 5×10^-9^ to claim significance as previously recommended for whole genome sequencing common variant association studies^14, 19^. The total numbers of variants that met our significance threshold were 2,214, 2,314, 2,697 and 2,442 for LDL-C, HDL-C, TC and TG, respectively, and after clumping^20^ the numbers of variants were 357, 338, 324, and 289, respectively. Of these variants, most were previously demonstrated to be associated with plasma lipids either at the variant- or locus-level^15^ (**Supplementary Table 3**, **Supplementary Fig. 2**).

To identify putative novel variant associations, we compared our results to a recent multi-ethnic lipid GWAS among 312,571 participants of the Million Veteran Program (MVP)^15^ as well as the GWAS Catalog (All associations(v1.0) file dated 06/04/2020) (**Fig. 2**). We clumped (window 250 kb, r^2^ 0.5) significant variants using Plink^20^ and queried these in the GWAS Catalog and MVP. Among genome-wide significant variants, we tabulated ‘known-position’ (variant previously associated), ‘known-loci’ (variants not previously significantly associated with the corresponding lipid phenotype but within 500 kb of a known locus, thereby representing additional allelic heterogeneity), and ‘novel’ variants (variants not in a known lipid locus) (**Supplementary Table 3**).

**Fig 2:**
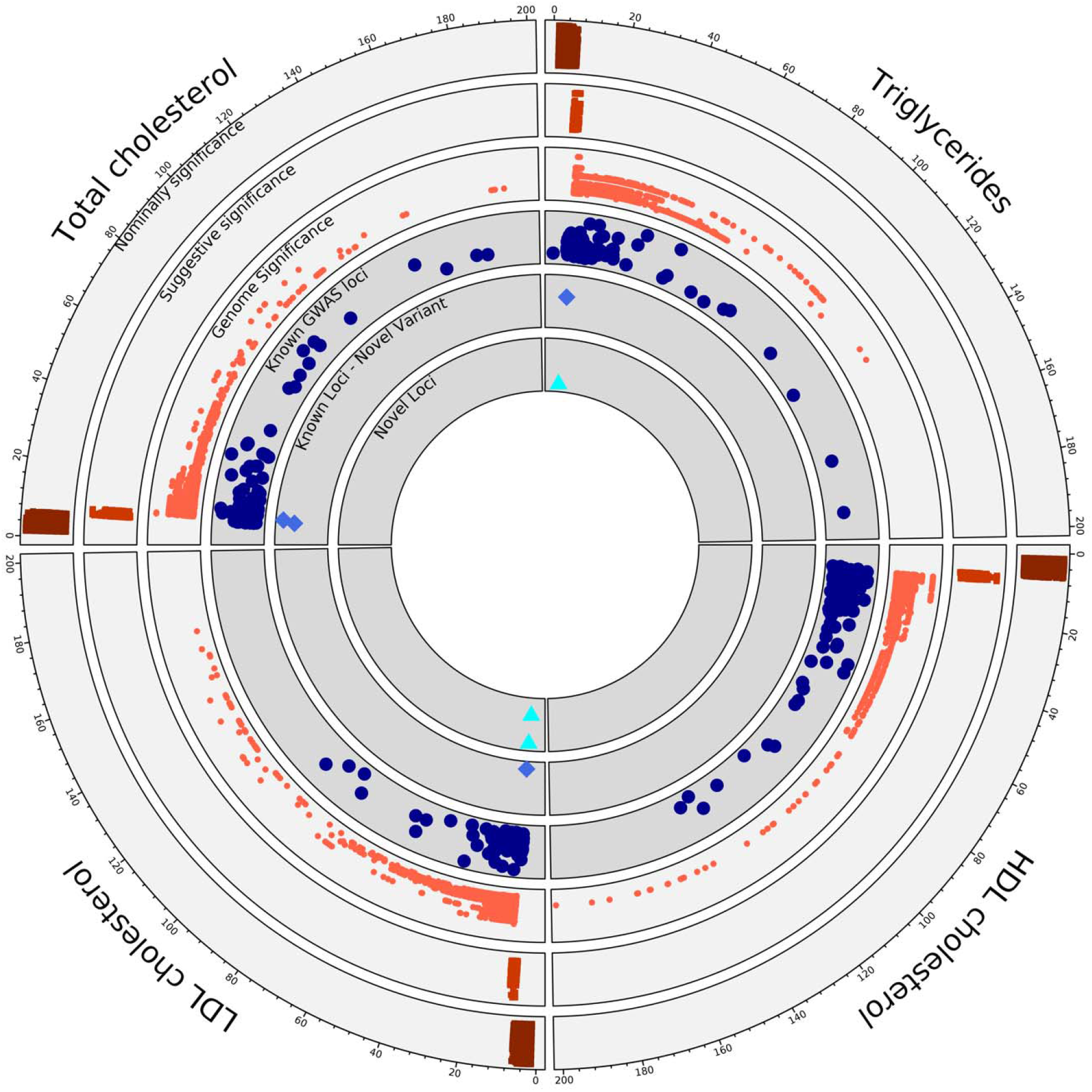
Summary of single variant genome wide association. Representation of the single variant GWAS results from TOPMed Freeze 8 whole genome sequenced data of 66,329 samples. Each quarter represents a different lipid phenotype, and dots extending in clock-wise fashion represent variants with increasing evidence of association as noted by -log10(p-value), which was truncated at 200. The outer three circles show the GWAS data from TOPMed freeze8 where variants binned to nominally significant (p-value 0.05 - 5×10^-07^), suggestive significant (p-value 5×10^-07^ - 5×10^-09^) and genome wide significant (p-value < 5×10^-09^). The inner three circles compare our TOPMed results with known significantly associated lipid loci and variants from the MVP summary statistics and GWAS catalog to the identified novel variants and loci that are genome-wide significant from the current study, respectively. TOPMed – Trans-Omics for Precision Medicine; GWAS – Genome Wide Association Study; MVP – Million Veteran Program.

The novel variants, tabulated in **Table 1**, are divided into two subsets – ‘novel variants’ or variants at established lipid loci for another lipid phenotype, and ‘novel loci,’ representing new loci associations for any lipid phenotype. For example, the *CETP* locus is well-known for its link to HDL-C, but we now found that rs183130 (16:56957451:C:T, MAF 28.3%) at the locus is associated with LDL-C. Similarly, the variants rs7140110 (13:113841051:T:C, MAF 27.8%) *GAS6* and rs73729083 (7:137875053:T:C, MAF 4.5%) *CREB3L2* are newly associated with TC, while previous studies showed that rs73729083 associates with LDL-C^21^ and rs7140110 associates with LDL-C^22^ and TG^23^. Index variants at novel loci were typically low frequency variants often observed in non-European ancestries, so we also conducted ancestry-specific association analyses for these alleles (**Supplementary Table 4**). For example, 12q23.1 (12:97352354:T:C, MAF 0.3%) and 4q34.2 (4:176382171:C:T, MAF 0.2%) associations with LDL-C are specific to Hispanic (MAF 1.3%) and Black (MAF 0.6%) populations, respectively and among Asians (MAF 1.5%) alone, 11q13.3 (11:69219641:C:T, MAF 0.2%) was associated with TG. One variant initially passing the novel locus filter for HDL-C (*RNF111* - rs112147665, beta = 8.664, p-value = 6.51×10^-10^), was in LD (r=0.7) with LIPC p.Thr405Met (rs113298164) which is known to be associated with HDL-C. The lead variant from MVP was 604 kb away from the *RNF111* variant but the rare *LIPC* missense variant p.Thr405Met was 421 kb away. Conditional analysis accounting for *LIPC* p.Thr405Met rendered the non-coding variant near *RNF111* variant non-significant (beta = 4.351, p-value = 2.47×10^-02^), therefore we reclassified *RNF111* variant as a known-position variant. Ancestry-specific GWAS did not yield additional novel loci beyond our larger trans-ancestry GWAS. Majority of genome significant single variants were captured by previous lipid GWAS^15^, but ancestry specific novel-hits are unique to WGS TOPMed data.

**Table 1.** Putative novel variants identified in TOPMed and evidence for replication. Variants identified as novel after comparing with the GWAS catalog and MVP summary statistics for associations with lipid phenotypes, including LDL-C, TC, and TG. All effect estimates are in mg/dL units, except for TG which was log-transformed in analysis thereby representing fractional change. Variants are categorized as novel loci or novel variant (i.e., known locus associated with another lipid phenotype) and the genes assigned to the variants per TOPMed whole genome sequence annotations (WGSA) are listed. Data is provided for the discovery (TOPMed freeze8) and replication cohorts (MGB Biobank and Penn Medicine Biobank). Meta-analysis with the replication cohorts was carried out and the corresponding beta and p-values are provided. GWAS – Genome Wide Association Study; MVP – Million Veteran Program; LDL-C – Low-Density Lipoprotein Cholesterol; TC – Total Cholesterol; TG – Triglycerides; TOPMed – Trans-Omics for Precision Medicine; WGSA – Whole Genome Sequence Annotations.

| Associated Lipid Phenotype | Novel variant class | Variants (Gene) | TOPMed Freeze8 (N=66329) |  |  | MGB Biobank (N=25137) |  |  | Penn Medicine Biobank (N=20079) |  |  | Meta Analysis (METASOFT) |  |
| --- | --- | --- | --- | --- | --- | --- | --- | --- | --- | --- | --- | --- | --- |
|  |  |  | TOPMed | TOPMed | TOPMed | MGB Biobank | MGB Biobank | MGB Biobank | Penn Medicine Biobank | Penn Medicine Biobank | Penn Medicine Biobank | Beta | P-value |
|  |  |  | Effect Estimate | P-value | MAF | Effect Estimate | P-value | MAF | Effect Estimate | P-value | MAF |  |  |
| LDL-C | Novel locus | 12:97352354:T:C | -12.439 | 4.88x10<sup>-09</sup> | 0.003 | 1.055 | 8.08x10<sup>-01</sup> | 0.002 | 11.441 | 3.19x10<sup>-01</sup> | 0.001 | 2.357 | 5.62 x10<sup>-01</sup> |
| LDL-C | Novel variant | 16:56957451:C:T<br>(CETP) | -1.568 | 2.88x10<sup>-09</sup> | 0.283 | -1.375 | 1.53x10<sup>-04</sup> | 0.309 | -2.35 | 1.54x10<sup>-04</sup> | 0.578 | -1.624 | 2.21 x10<sup>-07</sup> |
| LDL-C | Novel locus | 4:176382171:C:T | -16.086 | 2.82x10<sup>-09</sup> | 0.002 | -13.340 | 1.71x10<sup>-01</sup> | 0.001 | 4.716 | 3.52x10<sup>-01</sup> | 0.005 | 0.882 | 8.44 x10<sup>-01</sup> |
| TC | Novel variant | 13:113841051:T:C<br>(GAS6) | 1.731 | 1.12x10<sup>-09</sup> | 0.278 | 0.890 | 3.94x10<sup>-02</sup> | 0.304 | 0.416 | 5.50x10<sup>-01</sup> | 0.563 | 0.758 | 3.89 x10<sup>-02</sup> |
| TC | Novel variant | 7:137875053:T:C<br>(CREB3L2) | -4.106 | 7.54x10<sup>-11</sup> | 0.045 | -4.755 | 1.06x10<sup>-02</sup> | 0.013 | -3.365 | 7.62x10<sup>-03</sup> | 0.118 | -3.803 | 2.69 x10<sup>-04</sup> |
| TG | Novel locus | 11:69219641:C:T | 0.232 | 1.98x10<sup>-09</sup> | 0.002 | -0.047 | 7.33x10<sup>-01</sup> | 0.000 | 0.202 | 7.82x10<sup>-02</sup> | 0.001 | 0.101 | 2.53 x10<sup>-01</sup> |
| TG | Novel variant | 13:107551611:C:T<br>(FAM155A) | 0.052 | 6.78x10<sup>-10</sup> | 0.045 | 0.016 | 4.68x10<sup>-01</sup> | 0.014 | 0.039 | 3.26x10<sup>-01</sup> | 0.016 | 0.021 | 2.62 x10<sup>-01</sup> |

Due to the paucity of available diverse WGS datasets with lipids of comparable size, we pursued replication with two genome-wide array-based genotyped datasets imputed to TOPMed WGS^17, 24^ : Mass General Brigham (MGB) Biobank (N=25,137) and Penn Medicine Biobank (N=20,079)^25, 26^, the replication cohorts had diverse ancestry distribution, where non-European samples accounted for 15.77% in MGB Biobank and 51.20% in Penn Medicine Biobank (**Supplementary Table 5**). We brought seven putative novel variants with p-values < 5×10^-9^ forward for replication. The three common variants, rs183130 (*CETP*), rs7140110 (*GAS6*) and rs73729083 (*CREB3L2*), that were associated with both LDL-C and TC in TOPMed also replicated in MGB and two (rs183130, rs73729083) replicated in Penn Biobank at an alpha level of 0.05 and consistent direction of effect (**Table 1**). The two variants that were associated in both replication studies were most significantly associated among African Americans in TOPMed (rs183130: beta = −2.762 mg/dL, p-value = 5.71×10^-07^; rs73729083: beta = −3.725 mg/dL, p-value = 5.25×10^-07^). Low-frequency variants from specific ancestry groups associated with lipids in TOPMed were not replicated but we cannot rule out the possibility of reduced power due to general underrepresentation of non-white ancestry groups in the replication data. In exploratory analyses, we extended the same approach for variants discovered to have 5×10^-9^ < p-value < 5×10^-7^ but did not observe replication (**Supplementary Table 6**).

### *CETP* locus, HDL-C, and LDL-C

*CETP* is a well-recognized Mendelian HDL-C gene and the locus was previously known to be significantly associated with HDL-C, TC and TG at genome-wide significance^15^. Pharmacologic CETP inhibitors have shown strong associations with increased HDL-C but mixed effects for LDL-C reduction in clinical trials^27, 28, 29, 30^. We found that the *CETP* locus variant rs183130 (chr16:56957451:C:T, MAF 28.3%, intergenic variant) was associated with reduced LDL-C concentration (beta = −1.568 mg/dL, SE = 0.264, p-value = 2.88×10^-09^). The lead HDL-C-associated variant at the locus, rs3764261 (chr16:56959412:C:A, MAF 30.3%), was associated with 3.5 mg/dL increased HDL-C (p-value = 8.03×10^-283^), and rs183130 was associated with 3.9 mg/dL increased HDL-C (p-value < 1×10^-284^) as well. Among the ancestry groups analyzed, rs183130 was most significantly associated with LDL-C among those of African ancestry (beta = −2.762 mg/dL, p-value = 5.71×10^-07^) (**Supplementary Table 7**). We next investigated variants by their HDL-C and LDL-C effects within this locus (+/-500kb of rs183130 and rs3764261) (**Fig. 3**). We identified five variants showing at least suggestive (p-value < 5×10^-07^) association with both HDL-C and LDL-C. Though variants with strong LD (linkage disequilibrium) existed, ancestry-specific analyses showed that the stronger LDL-C effects were among those of African ancestry.

**Fig 3:**
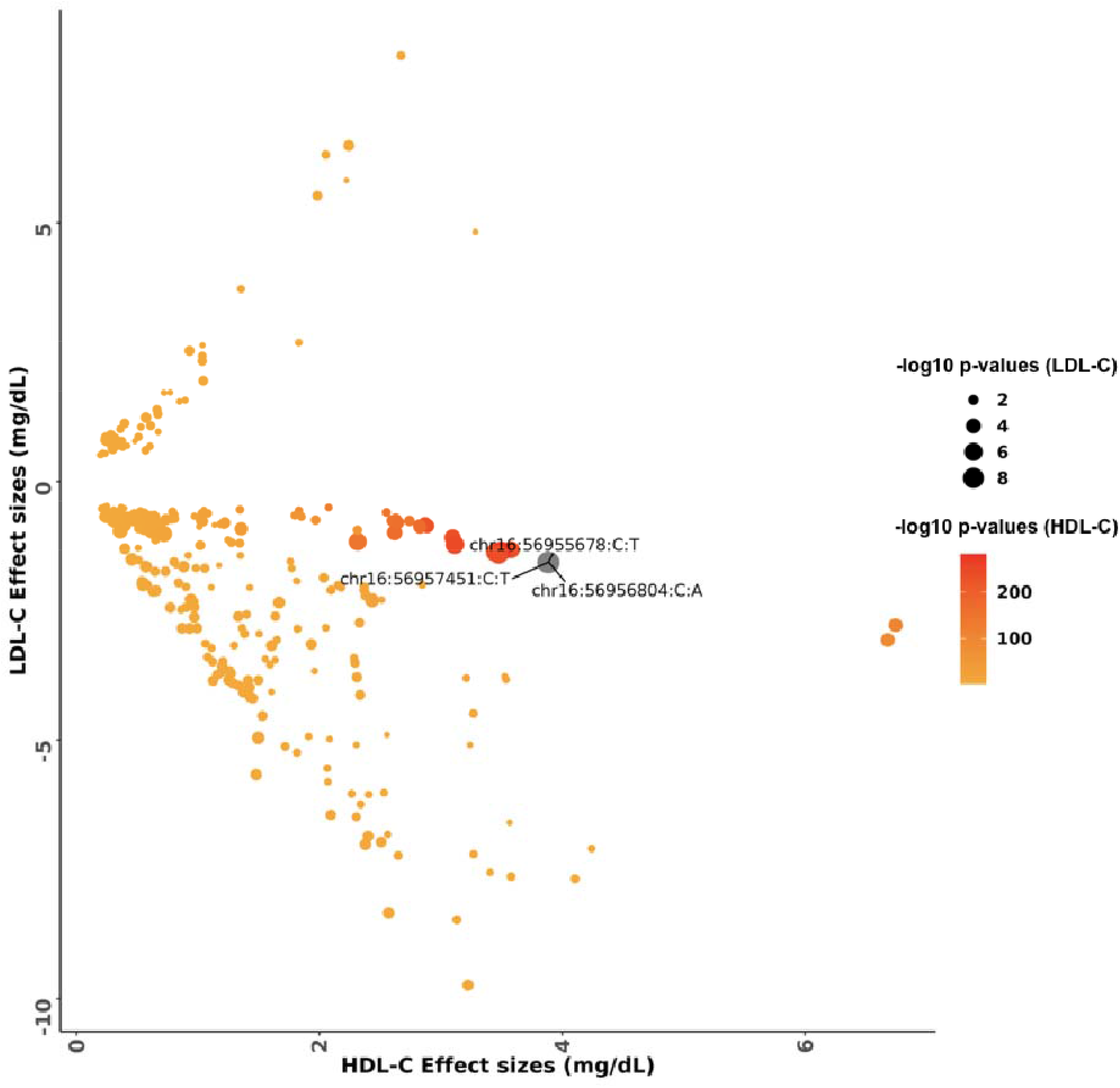
Comparison of effects estimates for HDL-C and LDL-C among variants in the *CETP* locus. The color scale of the data points was based on -log10 p-values from HDL-C association and the size of each data point was based on - log10 p-values of LDL-C association. Variants which are genome wide significant with LDL-C are represented as chromosome:position:reference allele:alternate allele. HDL-C – High-Density Lipoprotein Cholesterol; LDL-C – Low-Density Lipoprotein Cholesterol.

To better understand the mechanisms for HDL-C and LDL-C effects at the *CETP* locus, we pursued colocalization with eQTLs from 3 tissues (Liver, Adipose Subcutaneous and Adipose Visceral [Omentum]) from GTEx^31^. We analyzed 5 LDL-C and 441 HDL-C associated (p-values < 5×10^-07^) variants. We correlated eQTL effect estimates for genes at the locus with lipid outcome effect estimates. Indeed, *CETP* gene expression effects were strongly negatively correlated with HDL-C effects (Liver: ρ −0.933, p-value 4.01×10^-17^; Adipose Subcutaneous: ρ −0.762, p-value 8.87×10^-12^; Adipose Visceral: ρ −0.739, p-value 5.52×10^-10^) (**Supplementary Fig. 3**). However, *CETP* expression effects were not significantly correlated with LDL-C (Liver: ρ 0.007, p-value 0.99; Adipose Subcutaneous: ρ 0.344, p-value 0.57; Adipose Visceral: ρ - 0.59, p-value 0.29). Given the possibility that the observed lack of correlation for LDL-C could be due to reduced power from a limited number of variants attaining a suggestive p-value (< 5×10^-07^), we repeated the analysis with a subset of 122 nominally significant (p-value < 0.05) LDL-C associated variants in this locus. Indeed, *CETP* gene expression effects were strongly positively correlated with LDL-C effects (Liver: ρ 0.957, p-value 2.28×10^-08^; Adipose Subcutaneous: ρ 0.922, p-value 1.34 x10^-15^; Adipose Visceral: ρ 0.868, p-value 6.09×10^-11^).

### Rare variant aggregates associated with lipids

#### I) Gene-Centric associations

We next evaluated the association of aggregated rare (MAF<1%) variants, linked to protein-coding genes (‘gene-centric’). We employed a Bonferroni-corrected significance threshold of 0.05/20,000=2.5×10^-06^ for coding and non-coding gene-centric rare variant analyses (**Supplementary Fig. 4**). We identified 102 coding and 160 non-coding gene-centric rare variant aggregates significantly associated with at least one of the four plasma lipid phenotypes in nonconditional analysis (**Supplementary Table 8-9**). We secondarily conditioned our significant aggregate sets on variants individually associated with lipid levels from the GWAS catalog, MVP summary statistics and the TOPMed data. We identified 74 coding and 25 non-coding rare variants aggregates associated with at least one lipid level after conditional analyses (**Supplementary Table 10-11**).

Most of the coding gene-centric sets remained significant after secondary conditioning while a minority of non-coding gene-centric sets remained significant after conditioning. Significant genes identified from coding rare variant analyses included multiple known Mendelian lipid genes including *LCAT*, *LDLR*, and *APOB* (**Supplementary Table 10)**. *RFC2* putative loss-of-function mutations (combined allele frequency < 0.002%) were significantly associated with triglycerides (p-value 2×10^-06^) representing a putative novel association for triglycerides. The *RFC2* aggregate set (plof) was associated with reduced TG (beta = −0.89 for log[TG]). The persistently significant regions identified from non-coding rare variant analyses linked to genes included the UTR (untranslated region) for *CETP* and promoter-CAGE (CAGE-Cap Analysis of Gene Expression sites) around *APOA1* for HDL-C, and *APOE* promoter-CAGE, *APOE* enhancer-DHS (DHS - DNase hypersensitivity sites), and *EHD3* promoter-DHS for total cholesterol (**Supplementary Table 11)**. Most of the coding aggregates had larger effects compared to non-coding aggregates, and among the non-coding aggregates *SPC24* non-coding aggregate (enhancer-CAGE) at the *LDLR* locus had the strongest effect for LDL-C (beta = 2.320 mg/dL; p-value = 1.75×10^-05^).

#### II) Region-Based associations

We also performed unbiased region-based rare variant association analyses tiled across the genome with both static and dynamic window sizes. We first evaluated 2.6M regions statically at 2 kb size and 1 kb window overlap by the sliding window approach. Statistical significance was assigned at 0.05/(2.6×1^-06^)=1.88×10^-08^. We identified 28 significantly associated windows with at least one lipid phenotype. After conditioning on variants individually associated with the corresponding lipid phenotype, we identified two regions at *LDLR* still significantly associated with both total cholesterol and LDL-C although these regions included both intronic and exonic variants (**Supplementary Table 12**). *LDLR* intron 1, which encodes *LDLR-AS1* (LDLR antisense RNA 1) on the minus strand, had suggestive evidence for association with TC (p-value 3.17×10^-6^) with −2.76 mg/dL reduction in TC. A prior study identify that a common variant (rs6511720, MAF 0.11) in *LDLR* intron 1 is associated with increased *LDLR* expression in a luciferase assay and reduction in LDL-C^32^. When adjusting for rs6511720, the significance improved (p-value 1.43×10^−8^) with -3.35 mg/dL reduction in TC.

For dynamic window scanning of the genome, we implemented the SCANG method^33^. The SCANG procedure accounts for multiple testing by controlling the genome-wide error rate (GWER) at 0.1^33^. In the dynamic window-based workflow, STAAR-O detected 51 regions significantly associated with at least one lipid phenotype after conditioning on known variants (**Supplementary Table 13**). Most of the regions mapped to known Mendelian lipid genes, including *LCAT* (8.7×10^-13^) for HDL-C, and *LDLR*(2.4×10^-28^, 7.3×10^-26^) and *PCSK9* (2.9×10^-12^, 5.5×10^-12^) for LDL-C and TC, respectively. Exon 4 aggregates of *LDLR* were specifically associated with 20 mg/dL increase in LDL-C. *PCSK9* Exon2-Intron2 region spanning chr1:55043782-55045960 had significantly reduced LDL-C by 6 mg/dL (p-value = 3×10^-13^), and the effect persisted even with only Intron 2 rare variants of *PCSK9* (−5 mg/dL, p-value = 2×10^-8^). Strikingly, in secondary analyses, we found evidence for very large effects for rare variants in *LDLR* Introns 2 and 3 (+21 mg/dL, p-value = 7×10^-4^) and *LDLR* Introns 16 and 17 (+17 mg/dL, p-value = 0.02), similar to rare coding *LDLR* mutations. While 32 of the significant dynamic windows also included exonic regions, there were also several dynamic windows significantly independently associated with lipids not containing exonic regions. For example, four non-coding windows (two overlapping) at 2p24.1, which harbors the Mendelian *APOB* gene, were significantly associated with LDL-C. Intronic non-coding regions were associated with both LDL-C and TC -associated windows at *LPAL2*-*LPA*-*SLC22A3*; for example *LPAL2* Intron 3 was associated with a 3.7 mg/dL increase in TC. Non-coding TC-associated significant dynamic windows were near *TOMM40/APOE*. One rare variant signal observed was at *TOMM40* Intron 6, where the ‘poly-T’ variant in this region is on the *APOE4* haplotype and influences expressivity for Alzheimer’s disease age-of-onset^34,35^. For HDL-C, we identified significant non-coding windows at an intergenic region near *LPL* and *CD36* Intron 4. In the generation of the spontaneously hypertensive rat model, the deletion of intron 4 in *Cd36* with resultant *Cd36* deficiency has been mapped to defective fatty acid metabolism in this model^36^. Several regions significant in SCANG were not even nominally significant in burden association analyses indicating the likelihood of causal variants with bidirectional effects.

Several gene-centric non-coding aggregates associated with lipids near known monogenic lipid genes but mapped to another gene at the locus via annotations. Therefore, we performed downstream conditional analyses adjusting the gene-centric non-coding results for rare coding variants (MAF<1%) within known lipid monogenic genes (**Supplementary Table 14**). When accounting for both common and rare coding variants at the nearby familial hypercholesterolemia *LDLR* gene, *SPC24*-enhancer DHS was significantly associated with total cholesterol (p-value= 3.01×10^-11^) and with suggestive evidence for LDL-C (p-value= 1.57×10^-06^). In a similarly adjusted model, *LDLR*-enhancer-DHS showed a strong association with TC (p-value 5.18×10^-12^). When adjusting for known common variants as well as rare coding variants in *PCSK9*, both *PCSK9*-enhancer DHS and *PCSK9*-promoter DHS were significantly associated with total cholesterol. (**Fig. 4, Supplementary Fig. 5**). Through this procedure, *CETP* UTR retained significance for its independent association with HDL-C as well as the putatively novel gene *EHD3*-promoter DHS association with TC. However, the non-coding gene-centric *APOC3* and *APOE* associations were rendered non-significant for HDL-C and TC, respectively.

**Fig 4:**
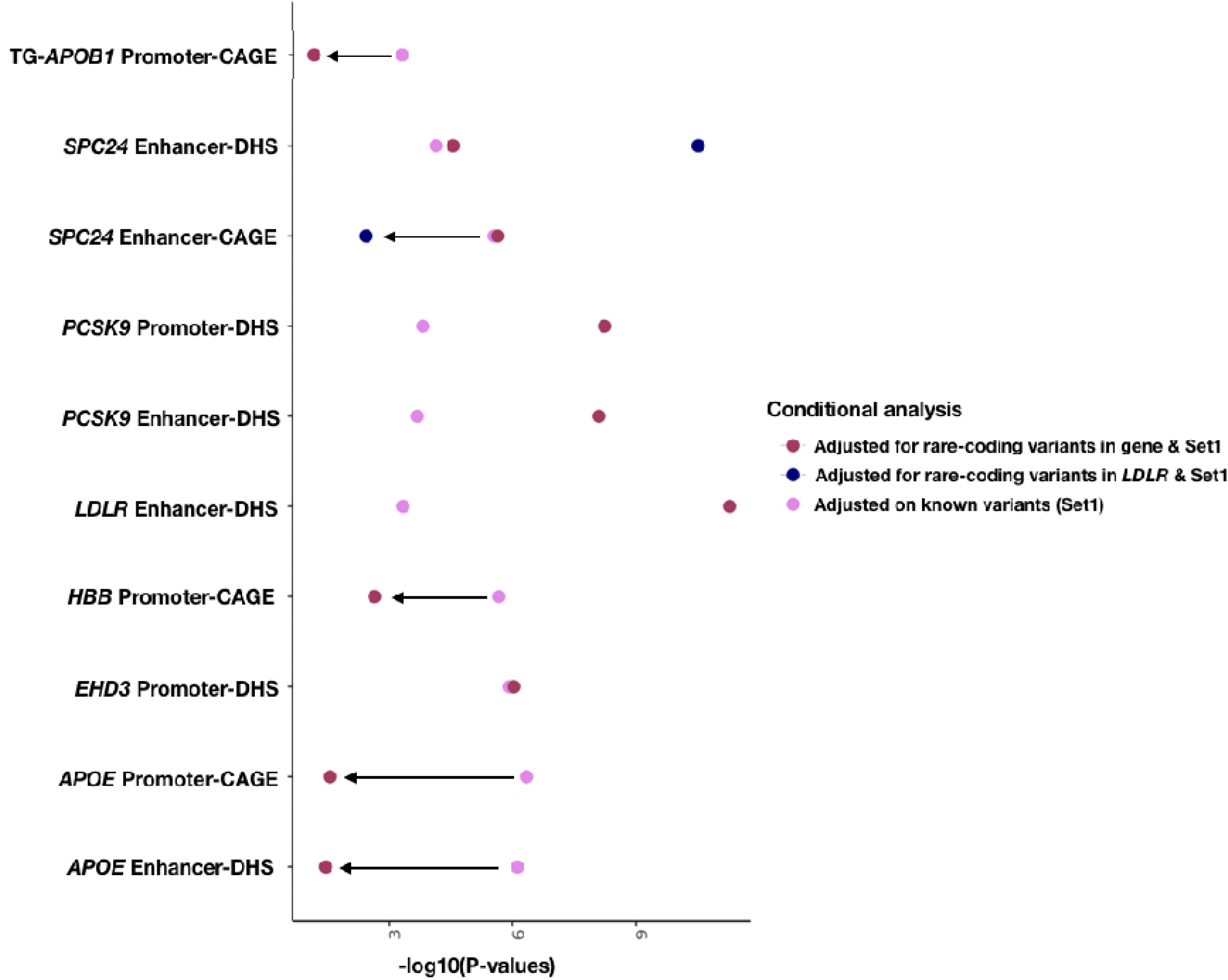
Conditional analysis of coding rare-variants from the same gene and a near-by gene. Non-coding rare variant sets significantly associated with TC and TG after the conditional analysis on known variants are shown with additional adjustment on rare-coding variants. The additional adjustment for rare-coding variants were carried out for the same gene of the aggregate set and for certain gene aggregates (*SPC24*) the conditional analysis was carried out with a nearby Mendelian gene. After adjusting for rare-coding variants and known variants, *EHD3* signal drops minimally, whereas signal from *PCSK9* (promoter-DHS, enhancer-DHS), *LDLR*-loci (enhancer-DHS, *SPC24* enhancer-DHS) enhances significantly. *APOB1*, *SPC24* (enhancer-CAGE), *HBB* and *APOE* signal drops after the conditional analysis on rare-coding variants. The different colored dots on the plot represents the conditional STAAR-O p-values when adjusting for known variants (Set1) and rare-coding variants of the same or near-by gene. STAAR – variant-Set Test for Association using Annotation information; TC – Total Cholesterol; TG – Triglycerides; CAGE – Cap Analysis of Gene Expression; DHS – DNase hypersensitivity.

Since we cannot rule out the possibility of reduced power for genome-wide rare variant analyses, we leveraged current knowledge of 22 Mendelian lipid genes for more focused exploratory analyses^14^. We validated most genes in rare variant coding analyses. The genes with the strongest coding signals typically had at least nominal evidence of gene-centric non-coding rare variant associations (**Supplementary Table 15, Supplementary Fig. 6**). When rare coding variants were introduced into the model, the evidence for non-coding rare variant associations were largely unchanged. Our findings expanding the currently described genetic basis for hypercholesterolemia to include rare non-coding variation at *LDLR* and *PCSK9* (**Fig. 5**).

**Fig 5:**
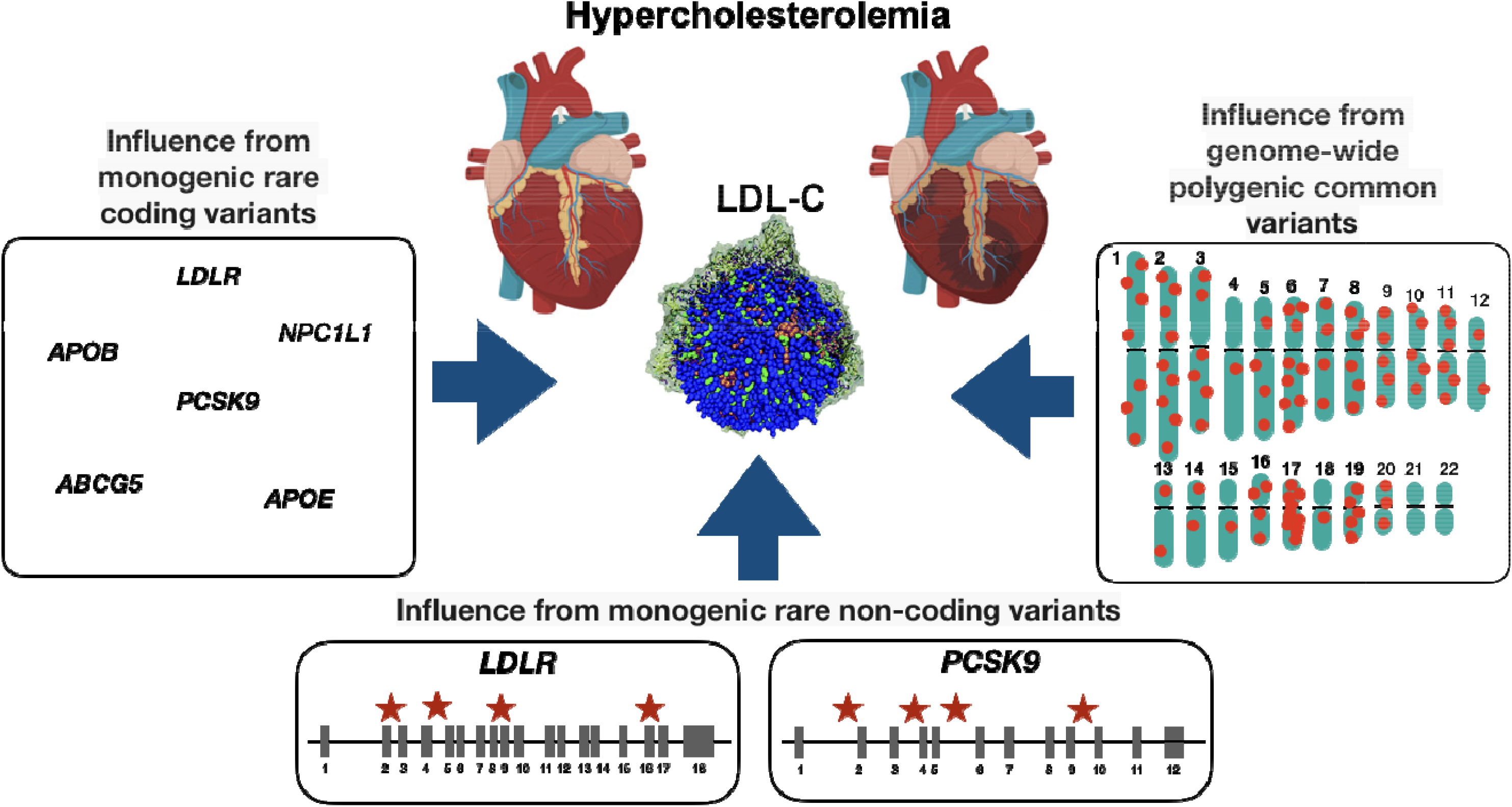
Influence of common and rare variants with hypercholesterolemia. In addition to monogenic contributions from rare variants in Mendelian hypercholesterolemia genes, multiple genome-wide significant LDL-C-associated common variants also yield a polygenic basis for hypercholesterolemia. In the present work, we now identify rare non-coding variants in proximity of Mendelian hypercholesterolemia genes, specifically *LDLR* and *PCSK9*, that also contribute to the genetic basis of hypercholesterolemia. LDL-C – Low-Density Lipoprotein Cholesterol

## Discussion

Conducting one of the largest population-based WGS association analyses, we now simultaneously interrogate and establish a common, rare coding, and rare non-coding variant model for a complex trait. Utilizing 66,329 diverse individuals with deep-coverage WGS, we interrogated 428M variants with plasma lipids expanding the allelic series to rare non-coding variants, often within introns, of Mendelian lipid genes with prior robust rare coding variant support. Our observations have important implications for plasma lipids as well as the genetic basis of complex traits more broadly.

WGS of diverse ancestries enables both allelic and locus heterogeneity for complex traits. Population genetic analyses have largely been enriched for individuals of European descent^37^. Genetic association of plasma lipids using arrays or whole exome sequencing among Europeans have yielded several important insights regarding plasma lipids and the causal determinants of CAD^5,4,38,39, 40^. Similar increasingly larger studies among non-Europeans have often yielded new genetic loci and sometimes new genes, such as *PCSK9*^1,15, 16, 41, 42^. Such differences have also led to concerns about the use of polygenic risk scores gleaned from much larger European GWAS of complex traits for non-Europeans^43^. Aided by the availability of WGS data, we identify new putative loci associated with lipids in non-Europeans. Furthermore, our study enabled the discovery of several novel alleles at known loci, with richly distinct allelic heterogeneity across ancestry groups. For example, HDL-C-raising *CETP* locus variants linked to *CETP* gene expression were only associated with LDL-C reduction among those of African ancestry. While all pharmacologic CETP inhibitors increase HDL-C, only those that decrease LDL-C also reduce cardiovascular disease risk^27, 28, 29, 30^. Given the contribution of genetic differences, clinical trials with more diverse samples would show insights.

Our study now provides increasingly robust evidence for a rare non-coding variant model for complex traits. Our rare non-coding variant associations in both gene-centric and sliding window models were largely restricted to the introns of Mendelian lipid genes with prior robust rare coding variant support consistent with biologic plausibility^44^. Rare intronic variants, often impacting splicing, have been previously implicated in afflicted Mendelian families or small exceptional case series, often through candidate gene approaches^45, 46, 47, 48^. We discovered one example of a rare non-coding signal without prior rare coding support – i.e., *EHD3*. We obtained estimates of phenotypic effect using burden tests. For most regions, even nominal significance was not detected using burden testing indicating the likelihood of variants with bidirectional effects further complicating clinical interpretation. When burden signals were detected, observed effects were typically larger than common non-coding variants and less than rare coding variants, with the exception of *LDLR*, consistent with whole genome mutational constraint models^49, 50, 51^.

The detection of independent rare non-coding variant signals has remained elusive largely due to limited sample sizes with requisite WGS and limitations in the interpretation of rare non-coding variation functional consequence. Previously, we used annotated functional non-coding sequence in 16,324 TOPMed participants, and found that rare non-coding gene regions associated with lipid levels, but they were not independent of individually associated single variants^14^. Using STAAR, we observed putative rare non-coding variant associations for lipids independent of individual variants associated with lipids in TOPMed.

WGS can improve diagnostic yield beyond the current standard of next-generation gene panel sequencing for dyslipidemias. A very small fraction with severe hypercholesterolemia and features consistent with strong genetic predisposition have a familial hypercholesterolemia variant in *LDLR*, *APOB*, or *PCSK9*^52, 53^. The presence of familial hypercholesterolemia variants is independently prognostic for CAD, beyond lipids, and merits the consideration of more costly lipid-lowering medications^52, 51, 55^. We now observe that rare *LDLR* variants in Introns 2, 3, 16, and 17 lead to approximately 0.5 standard deviation increase in LDL-C, approximating effects observed with clinically reported exonic familial hypercholesterolemia variants in *LDLR*^55^. Small studies have indicated the possibility of rare intronic *LDLR* variants causing familial hypercholesterolemia due to altered splicing, which we now observe in our unbiased population-based WGS study^56, 57^. A WGS approach to lipid disorders, particularly for familial hypercholesterolemia, will markedly improve the diagnostic yield beyond existing limited approaches.

Our dynamic window approach may also improve the clinical curation of exonic variants. Among the data used to curate exonic variants is the use of *in silico* functional prediction tools^58^. Although evolutionary constraint measures are typically employed, such tools are largely agnostic to functional domain. As it relates to lipids, disruptive *APOB* and *PCSK9* exonic variants can lead to strikingly opposing directions with large effects for LDL-C depending on locations^1,8,59, 60^. Using SCANG^33^, we detect a significant association with large effect for *LDLR* Exon 4 itself. This observation supports the pathogenicity of *LDLR* Exon 4 disruptive variants among patients with severe hypercholesterolemia. The majority of familial hypercholesterolemia variants worldwide occur in Exon 4 of *LDLR*^61, 62, 63, 64^. Conventional rare coding variant analyses aggregate all exonic variants for a transcript. Here, we demonstrate an opportunity for exon-level rare variant association testing.

Our study has important limitations. First, while our study is large for a WGS study by contemporary standards, it is dwarfed by existing GWAS datasets limiting power for novel discovery. Nevertheless, by using WGS in diverse ancestries, we can study hundreds of millions new variants. Second, prediction of rare non-coding variation consequence to prioritize causal variants remains a challenge thereby limiting power^65^. The striking difference for most STAAR and burden results also highlights bidirectional effects for rare non-coding variants within the same region and further challenges for clinical utility. Third, given the paucity of multi-ancestral WGS datasets with lipids, our analyses are largely restricted to TOPMed. For single variant associations, we pursued TOPMed-imputed GWAS datasets but were limited by the lack of ancestral diversity. As TOPMed is a consortium of multiple different cohorts, we demonstrate consistencies by cohort. Furthermore, rare variant non-coding signals were largely restricted to regions with rare variant coding signals supporting biological plausibility.

In conclusion, using WGS and lipids among 66,329 ancestrally diverse individuals we expand the catalog of alleles associated with lipids, including allelic heterogeneity at known loci and locus heterogeneity by ancestry. We characterize the common, rare coding, and rare non-coding variant model for lipids. Lastly, we now demonstrate a monogenic-equivalent model for rare *LDLR* intronic variants predisposing to marked alterations in LDL-C, currently not recognized in current population or clinical models for LDL-C.

### Online Methods Dataset

#### i) Contributing studies

The discovery cohort includes whole genome sequenced (WGS) data of 66,329 samples from 21 studies of the Trans-Omics for Precision Medicine (TOPMed) program with blood lipids available^17^. The overall goal of TOPMed is to generate and use trans-omics, including whole genome sequencing, of large numbers of individuals from diverse ancestral backgrounds with rich phenotypic data to gain novel insights into heart, lung, blood, and sleep disorders. The Freeze 8 data includes 140,306 samples out of which 66,329 samples qualified with lipid phenotype. Freeze 8 dataset passed the central quality control protocol implemented by the TOPMed Informatics Research Core (described below) and was deposited in the dbGaP TOPMed Exchange Area.

The studies included in the current dataset, along with their abbreviations and sample sizes, contains the Old Order Amish (Amish, n=1,083), Atherosclerosis Risk in Communities study (ARIC, n=8,016), Mt Sinai BioMe Biobank (BioMe, n=9,848), Coronary Artery Risk Development in Young Adults (CARDIA, n=3,056), Cleveland Family Study (CFS, n=579), Cardiovascular Health Study (CHS, n=3,456), Diabetes Heart Study (DHS, n=365), Framingham Heart Study (FHS, n=3,992), Genetic Studies of Atherosclerosis Risk (GeneSTAR, n=1,757), Genetic Epidemiology Network of Arteriopathy (GENOA, n=1,046), Genetic Epidemiology Network of Salt Sensitivity (GenSalt, n=1,772), Genetics of Lipid-Lowering Drugs and Diet Network (GOLDN, n=926), Hispanic Community Health Study - Study of Latinos (HCHS_SOL, n=7714), Hypertension Genetic Epidemiology Network and Genetic Epidemiology Network of Arteriopathy (HyperGEN, n=1,853), Jackson Heart Study (JHS, n=2,847), Multi-Ethnic Study of Atherosclerosis (MESA, n=5,290), Massachusetts General Hospital Atrial Fibrillation Study (MGH_AF, n=683), San Antonio Family Study (SAFS, n=619), Samoan Adiposity Study (SAS, n=1,182), Taiwan Study of Hypertension using Rare Variants (THRV, n=1,982) and Women’s Health Initiative (WHI, n=8,263) (Please see **Supplementary Text**for additional details). The multi-ancestral data set included individuals from White (44%), Black (26%), Hispanic (21%), Asian (7%), and Samoan (2%) ancestries. Study participants granted consent per each study’s Institutional Review Board (IRB) approved protocol. Secondarily, these data were analyzed through a protocol approved by the Massachusetts General Hospital IRB. **Supplementary Table 1** details the number of samples across different studies and ancestral group.

The replication cohorts include TOPMed-imputed genome-wide array data from the Mass General Brigham (MGB) and Penn Medicine Biobanks which consist of 25,137 samples and 20,079 samples respectively^25, 26^. We curated the MGB Biobank and Penn Medicine Biobank phenotype data from the corresponding electronic health record databases in accordance with corresponding institutional IRB approvals. Consent was previously obtained from each participant regarding storage of biological specimens, genetic sequencing, access to all available electronic health record (EHR) data, and permission to recontact for future studies. The MGB Biobank consists of 54% and Penn Medicine Biobank consist of 52% female samples and average ages were 55.89 years and 58.35 years, respectively (**Supplementary Table 5**).

#### ii) Phenotypes

The primary outcomes in this study included LDL cholesterol (LDL-C), HDL cholesterol (HDL-C), total cholesterol (TC) and triglycerides (TG) phenotypes. LDL-C was either directly measured or calculated by the Friedewald equation when triglycerides were <400 mg/dL. Given the average effect of lipid lowering-medicines, when lipid-lowering medicines were present, we adjusted the total cholesterol by dividing by 0.8 and LDL-C by dividing by 0.7, as previously done^14^. Triglycerides remained natural log transformed for analysis. Fasting status was accounted for with an indicator variable.

We harmonized the phenotypes across each cohort^18^ and inverse rank normalization of the residuals of each race within each cohort scaled by the standard deviation of the trait and adjusted for covariates^12^. We included covariates such as age, age^2^, sex, PC1-11, study-groups as well as Mendelian founder lipid variants *APOB* p.R3527Q and *APOC3* p.R19X for the Amish cohort^7,66,8^. **Supplementary Table 1** provides the distributions of each of the four lipid phenotypes by cohort, ancestral groups, and gender. We executed similar steps of phenotype harmonization and normalization for the replication cohorts. Additionally, we adjusted the MGB Biobank for study-center and array-type, and Penn Medicine Biobank for ancestry and BMI in addition to the other common covariates.

#### iii) Genotypes

Whole genome sequencing of goal >30X coverage was performed at seven centers (Broad Institute of MIT and Harvard, Northwest Genomics Center, New York Genome Center, Illumina Genomic Services, PSOMAGEN [formerly Macrogen], Baylor College of Medicine Human Genome Sequencing Center and McDonnell Genome Institute [MGI] at Washington University). In most cases, all samples for a given study within a given Phase were sequenced at the same center (**Supplementary Text**). The reads were aligned to human genome build GRCh38 using a common pipeline across all centers (BWA-MEM).

The TOPMed Informatics Research Core at the University of Michigan performed joint genotype calling on all samples in Freeze 8. The variant calling “GotCloud” pipeline (https://github.com/statgen/topmed_variant_calling) is under continuous development and details on each step can be accessed through TOPMed website for Freeze8 (https://www.nhlbiwgs.org/topmed-whole-genome-sequencing-methods-freeze-8)17. The resulting BCF files were split by study and consent group for distribution to approved dbGaP users. Quality control was performed at each stage of the process, poor variant quality was indicated by missing rate >20%, mappability score <0.8, mean depth of coverage >500X, and Ti/Tv ratio, by the Sequencing Centers, the IRC and the TOPMed Data Coordinating Center (DCC). The VCF/BCF files were converted to GDS (Genomic Data Structure) format by the DCC and were deposited into the dbGap TOPMed Exchange Area.

The genetic relationship matrix (GRM) is an N*N matrix of relatedness information of the samples included in the study and was computed centrally using ‘PC-relate’ R package (version: 1.24.0)^67^. Using the ‘Genesis’ R package (version:2.20.1)^68^ we generated subsetted GRM for the samples with plasma lipid profiles. The GDS files with the variants were annotated internally by curating data from multiple database sources using Functional Annotation of Variant–Online Resource (FAVOR (http://favor.genohub.org)^13^. This study used the resultant aGDS (annotation GDS) files.

The MGB Biobank replication cohort was genotyped using three different arrays (Multiethnic Exome Global (Meg), Human multi-ethnic array (Mega), and Expanded multi-ethnic genotyping array (Megex)), and we separately imputed the data using TOPMed imputation server with default parameters^69, 70^. This study applied the Version-r2 of the imputation panel, it includes 97,256 reference samples and ∼300M genetic variants. The Illumina Global Screening array was used to genotype the Penn Medicine Biobank. Penn Medicine Biobank TOPMed imputation was performed using EAGLE^70^ and Minimac^71^ software. For this study we downloaded variants that passed a min R^2^ threshold of 0.3.

### Single Variant Association

We performed genome-wide single variant association analyses for autosomal variants with minor allele frequency (MAF) greater than 0.1% across the dataset with each of the four lipid phenotypes. We implemented the SAIGE-QT^72^ method, which employs fast linear mixed models with kinship adjustment, in Encore (https://encore.sph.umich.edu/) for single variant association analyses. We additionally adjusted the model for covariates (PC1-PC11, age, sex, age^2^, and study-groups [cohort-race subgrouping]).

We conducted single variant association replications for putative novel variants. After comparing the results with published lipid GWAS summary statistics, we filtered putative novel GWAS variants based on a stringent whole genome-wide significant threshold (alpha = 5×10^-9^)^73^. Replication was performed in the MGB and Penn Medicine Biobanks where models were fitted as indicated above. Additionally, we adjusted the MGB Biobank for study recruitment center and array and Penn Medicine Biobank for ancestry and BMI. In the MGB Biobank, we selected lipid concentrations closest to the sample acquisition time point and adjusted for statins if prescribed within one year prior to sample acquisition. In the Penn Biobank, we utilized each participant’s median lipid concentration for replication; statins prescribed prior to lipid concentration used were adjusted in the models. Additionally, we carried out meta-analysis using fixed effects model based on inverse-variance-weighted effect size for the two replication cohorts using METASOFT^74^.

### Rare variant association test

We performed rare variant association (RVA) using the Variant-Set Test for Association using Annotation infoRmation (STAAR) pipeline^13^ from STARtopmed R package. STAARpipeline is a regression-based framework that permits adjustment of covariates, population structure, and relatedness by fitting linear and logistic mixed models for quantitative and dichotomous traits^75^,. We chose STAAR to leverage the annotation information and associated scores that were available for TOPMed Freeze 8 data to incorporate the analysis of rare non-coding variants from whole genome sequencing. The method implements genome-wide scanning of rare variants (MAF<0.01) in gene-centric and region-based workflows. For each variant set, STAARpipeline calculates a set-based p-value using the STAAR method, which increases the analysis power by incorporating multiple *in silico* variant functional annotation scores capturing diverse genomic features and biochemical readouts^13^. We aggregated rare variants into multiple groups for coding and non-coding analyses. For the coding region, we defined five different aggregate masks of rare variants 1) plof (putative loss-of-function), plof-Ds (putative loss-of-function or disruptive missense), missense, disruptive-missense, and synonymous. For the non-coding regions, we used seven rare variant masks: 1) promoter-CAGE (promoter variants within Cap Analysis of Gene Expression [CAGE] sites^77, 78^), 2) promoter-DHS (promoter variants within DNase hypersensitivity [DHS] sites ^79^), 3) enhancer-CAGE (enhancer within CAGE sites^78^), 4) enhancer-DHS (enhancer variants within DHS sites^80^), 5) UTR (rare variants in 3’ untranslated region [UTR] and 5’ UTR untranslated region), 6) upstream, and 7) downstream. Detailed explanations of the regions defined based on these masks is discussed within STAARpipeline^13^.

In the gene-centric workflows, for both coding (within exonic boundaries) and non-coding (promoter: +/-3kb window of transcription starting site (TSS), enhancer: GeneHancer predicted regions) regions, we considered only genes with at least two rare variants (i.e., 18,445 genes in all 22 autosomes). In the region-based workflows, we implemented two protocols: 1) a ‘sliding window’ approach, where we aggregated rare variants within 2-kb sliding windows and with 1-kb overlap length, and 2) a ‘dynamic window’ approach, where we executed SCANG^33^ method and aggregated dynamically variant-sets between 40-300 variants per set, where the method systematically scans the whole genome with overlapping windows of varying sizes. The STAARtopmed R-package implements multiple rare-variants aggregate tests including SKAT^81^, Burden^82^ and ACAT^83^ and integrates them as STAAR-O^13^. We performed gene-centric and region-based rare variant tests using annotated GDS files of TOPMed.

We completed aggregate tests as three-step process. In the first step, we fitted a null model using glmmkin() function in STAARtopmed. The null model was fitted for each of the four lipid phenotypes adjusted for all covariates and relatedness except the genotype of interest. In the second step, we ran genome wide gene-centric and region-based rare-variant aggregate tests. The third step directed conditional analyses, where the results were adjusted for previously known significantly lipid-associated (i.e., p < 5×10^-8^ in external datasets) individual variants from GWAS Catalog^84^ and Million Veterans Program (MVP)^15^ GWAS summary statistics. To obtain effect estimates of significant aggregate sets, we associated the cumulative genotypes (binary scores) based on the variants forming the aggregates and used Glmm.Wald test from GMMAT R package^75^(version 1.3.1). For significantly-associated window-based rare variant aggregations, we trimmed the exonic variants and estimated the effects with only non-coding variants.

### *CETP* gene expression and lipid trait colocalization

We studied the correlation of LDL-C and HDL-C effects with eQTL effects at chromosome 16q13, which includes *CETP*. We downloaded GTEx eQTL build 38 (version8) data for Liver, Adipose Subcutaneous and Adipose Visceral (Omentum) tissues from GTEx Portal on 16/APR/2020^85^. We selected eQTLs with nominal significance (p-value<0.05) and utilized the eQTL-gene pairs with the most significant p-values. Genes with at least 5 eQTLs were selected for the colocalization analysis. We selected variants with a suggestive significance (p-value < 5×10^-7^) for LDL-C or HDL-C effects within 500 kb of the lead locus variant. We performed Pearson correlation tests between the lipid effect estimates and gene expression effects (slope) from GTEX.

## Acknowledgments

Whole genome sequencing (WGS) for the Trans-Omics in Precision Medicine (TOPMed) program was supported by the National Heart, Lung and Blood Institute (NHLBI). P.N. is supported by grants from the National Heart, Lung, and Blood Institute (R01HL142711, R01HL148050, R01HL151283, R01HL148565, R01HL135242, R01HL151152), Fondation Leducq (TNE-18CVD04), and Massachusetts General Hospital (Paul and Phyllis Fireman Endowed Chair in Vascular Medicine). The Amish studies were supported by NIH grants R01 AG18728, U01 HL072515, R01 HL088119, R01 HL121007, and P30 DK072488. The Atherosclerosis Risk in Communities (ARIC) study has been funded in whole or in part with Federal funds from the National Heart, Lung, and Blood Institute, National Institutes of Health, Department of Health and Human Services (contract numbers HHSN268201700001I, HHSN268201700002I, HHSN268201700003I, HHSN268201700004I and HHSN268201700005I). The authors thank the staff and participants of the ARIC study for their important contributions. The Mount Sinai BioMe Biobank (BioMe) has been supported by The Andrea and Charles Bronfman Philanthropies and in part by Federal funds from the NHLBI and NHGRI (U01HG00638001; U01HG007417; X01HL134588). Coronary Artery Risk Development in Young Adults (CARDIA) Study (phs001612) was performed at the Baylor College of Medicine Human genome Sequencing Center (contract HHSN268201600033I). Core support including centralized genomic read mapping and genotype calling, along with variant quality metrics and filtering were provided by the TOPMed Informatics Research Center (3R01HL-117626-02S1; contract HHSN268201800002I). Core support including phenotype harmonization, data management, sample-identity QC, and general program coordination were provided by the TOPMed Data Coordinating Center (R01HL-120393; U01HL-120393; contract HHSN268201800001I). We gratefully acknowledge the studies and participants who provided biological samples and data for TOPMed. The Coronary Artery Risk Development in Young Adults Study (CARDIA) is conducted and supported by the National Heart, Lung, and Blood Institute (NHLBI) in collaboration with the University of Alabama at Birmingham (HHSN268201800005I & HHSN268201800007I), Northwestern University (HHSN268201800003I), University of Minnesota (HHSN268201800006I), and Kaiser Foundation Research Institute (HHSN268201800004I). Cleveland Family Study (CFS) is supported by grants from the NHLBI (HL046389, HL113338, and 1R35HL135818). Cardiovascular Health Study (CHS) was supported by contracts HHSN268201200036C, HHSN268200800007C, HHSN268201800001C, N01HC55222, N01HC85079, N01HC85080, N01HC85081, N01HC85082, N01HC85083, N01HC85086, 75N92021D00006, and grants U01HL080295 and U01HL130114 from the National Heart, Lung, and Blood Institute (NHLBI), with additional contribution from the National Institute of Neurological Disorders and Stroke (NINDS). Additional support was provided by R01AG023629 from the National Institute on Aging (NIA). A full list of principal CHS investigators and institutions can be found at CHS-NHLBI.org. The content is solely the responsibility of the authors and does not necessarily represent the official views of the National Institutes of Health. Diabetes Heart Study (DHS) was supported by HL92301, HL67348, NS058700, AR48797, DK071891, AG058921, the General Clinical Research Center of the Wake Forest University School of Medicine (RR07122, HL085989), the American Diabetes Association, and a pilot grant from the Claude Pepper Older Americans Independence Center of Wake Forest University Health Sciences (AG10484). Framingham Heart Study (FHS) acknowledges the support of contracts NO1-HC-25195 and HHSN268201500001I from the National Heart, Lung and Blood Institute and grant supplement R01 HL092577-06S1 for this research. WGS for “NHLBI TOPMed: Whole Genome Sequencing and Related Phenotypes in the Framingham Heart Study” (phs000974) was performed at the Broad Institute of MIT and Harvard (HHSN268201500014C, 3R01HL092577-06S1, and 3U54HG003067-12S2). We also acknowledge the dedication of the FHS study participants without whom this research would not be possible. Genetic Studies of Atherosclerosis Risk (GeneSTAR) was supported by grants from the National Institutes of Health/National Heart, Lung, and Blood Institute (U01 HL72518, HL087698, HL49762, HL58625, HL071025, HL112064), the National Institutes of Health/National Institute of Nursing Research (NR0224103), and by a grant from the National Institutes of Health/National Center for Research Resources (M01-RR000052) to the Johns Hopkins General Clinical Research Center. Genetic Epidemiology Network of Arteriopathy (GENOA) was supported by the National Heart, Lung and Blood Institute (HL054457, HL054464, HL054481, HL087660, and HL119443) of the National Institutes of Health. Genetic Epidemiology Network of Salt Sensitivity (GenSalt) was supported by research grants (U01HL072507, R01HL087263, and R01HL090682) from the National Heart, Lung and Blood Institute, National Institutes of Health, Bethesda, MD. Genetics of Lipid-Lowering Drugs and Diet Network (GOLDN) biospecimens, baseline phenotype data, and intervention phenotype data were collected with funding from National Heart, Lung and Blood Institute (NHLBI) grant U01 HL072524. The Hispanic Community Health Study/Study of Latinos (HCHS-SOL) was carried out as a collaborative study supported by contracts from the National Heart, Lung, and Blood Institute (NHLBI) to the University of North Carolina (N01-HC65233), University of Miami (N01-HC65234), Albert Einstein College of Medicine (N01-HC65235), Northwestern University (N01-HC65236), and San Diego State University (N01-HC65237). The Hypertension Genetic Epidemiology Network and Genetic Epidemiology Network of Arteriopathy (HyperGEN) Study is part of the National Heart, Lung, and Blood Institute (NHLBI) Family Blood Pressure Program; collection of the data represented here was supported by grants U01 HL054472 (MN Lab), U01 HL054473 (DCC), U01 HL054495 (AL FC), and U01 HL054509 (NC FC). The HyperGEN: Genetics of Left Ventricular Hypertrophy Study was supported by NHLBI grant R01 HL055673 with whole-genome sequencing made possible by supplement −18S1. The Jackson Heart Study (JHS) is supported and conducted in collaboration with Jackson State University (HHSN268201800013I), Tougaloo College (HHSN268201800014I), the Mississippi State Department of Health (HHSN268201800015I) and the University of Mississippi Medical Center (HHSN268201800010I, HHSN268201800011I and HHSN268201800012I) contracts from the National Heart, Lung, and Blood Institute (NHLBI) and the National Institute on Minority Health and Health Disparities (NIMHD). The authors also wish to thank the staffs and participants of the JHS. Multi-Ethnic Study of Atherosclerosis (MESA) and the MESA SHARe projects are conducted and supported by the National Heart, Lung, and Blood Institute (NHLBI) in collaboration with MESA investigators. Support for MESA is provided by contracts 75N92020D00001, HHSN268201500003I, N01-HC-95159, 75N92020D00005, N01-HC-95160, 75N92020D00002, N01-HC-95161, 75N92020D00003, N01-HC-95162, 75N92020D00006, N01-HC-95163, 75N92020D00004, N01-HC-95164, 75N92020D00007, N01-HC-95165, N01-HC-95166, N01-HC-95167, N01-HC-95168, N01-HC-95169, UL1-TR-000040, UL1-TR-001079, and UL1-TR-001420. Funding for SHARe genotyping was provided by NHLBI Contract N02-HL-64278. Genotyping was performed at Affymetrix (Santa Clara, California, USA) and the Broad Institute of Harvard and MIT (Boston, Massachusetts, USA) using the Affymetrix Genome-Wide Human SNP Array 6.0. Also supported in part by NHLBI CHARGE Consortium Contract HL105756. The provision of genotyping data was supported in part by the National Center for Advancing Translational Sciences, CTSI grant UL1TR001881, and the National Institute of Diabetes and Digestive and Kidney Disease Diabetes Research Center (DRC) grant DK063491 to the Southern California Diabetes Endocrinology Research Center. Infrastructure for the CHARGE Consortium is supported in part by the National Heart, Lung, and Blood Institute (NHLBI) grant R01HL105756. The Massachusetts General Hospital Atrial Fibrillation Study (MGH-AF) was supported by grants to Dr. Ellinor from the Fondation Leducq (14CVD01), the National Institutes of Health to Dr. Ellinor (1RO1HL092577, R01HL128914, K24HL105780) and Dr. Lubitz (1R01HL139731) and by grants from the American Heart Association to Dr. Ellinor (18SFRN34110082) and to Dr. Lubitz (18SFRN34250007). San Antonio Family Study (SAFS) was supported in part by National Institutes of Health (NIH) grants R01 HL045522, MH078143, MH078111 and MH083824; and whole genome sequencing of SAFS subjects was supported by U01 DK085524 and R01 HL113323. We are very grateful to the participants of the San Antonio Family Study for their continued involvement in our research programs. Samoan Adiposity Study (SAS) was funded by NIH grant R01-HL093093. We thank the Samoan participants of the study and local village authorities. We acknowledge the support of the Samoan Ministry of Health and the Samoa Bureau of Statistics for their support of this research. The Rare Variants for Hypertension in Taiwan Chinese (THRV) is supported by the National Heart, Lung, and Blood Institute (NHLBI) grant (R01HL111249) and its participation in TOPMed is supported by an NHLBI supplement (R01HL111249-04S1). SAPPHIRe was supported by NHLBI grants (U01HL54527, U01HL54498) and Taiwan funds, and the other cohorts were supported by Taiwan funds. The Women’s Health Initiative (WHI) program is funded by the National Heart, Lung, and Blood Institute, National Institutes of Health, U.S. Department of Health and Human Services through contracts HHSN268201600018C, HHSN268201600001C, HHSN268201600002C, HHSN268201600003C, and HHSN268201600004C. The Centers for Common Disease Genomics (CCDG) program was supported by NHGRI and NHLBI, and whole genome sequencing was performed at the Baylor College of Medicine Human Genome Sequencing Center (UM1 HG008898 and R01HL059367). We like to acknowledge all the grants that supported this study, R01 HL121007, U01 HL072515, R01 AG18728, X01HL134588, HL 046389, HL113338, and 1R35HL135818, K01 HL135405, R03 HL154284, U01HL072507, R01HL087263, R01HL090682, P01HL045522, R01MH078143, R01MH078111, R01MH083824, U01DK085524, R01HL113323, R01HL093093, R01HL140570, R01HL142711, R01HL127564, R01HL148050, R01HL148565, and Leducq TNE-18CVD04. The views expressed in this manuscript are those of the authors and do not necessarily represent the views of the National Heart, Lung, andBlood Institute; the National Institutes of Health; or the U.S.Department of Health and Human Services.

## Author’s information

### NHLBI Trans-Omics for Precision Medicine (TOPMed) Consortium

Namiko Abe^63^, Gonçalo Abecasis^64^, Francois Aguet^65^, Christine Albert^66^, Laura Almasy^67^, Alvaro Alonso^68^, Seth Ament^69^, Peter Anderson^70^, Pramod Anugu^71^, Deborah Applebaum-Bowden^72^, Kristin Ardlie^65^, Dan Arking^73^, Allison Ashley-Koch^74^, Tim Assimes^75^, Paul Auer^76^, Dimitrios Avramopoulos^73^, Najib Ayas^77^, Adithya Balasubramanian^78^, John Barnard^79^, Kathleen Barnes^80^, R. Graham Barr^81^, Emily Barron-Casella^73^, Lucas Barwick^82^, Terri Beaty^73^, Gerald Beck^83^, Diane Becker^84^, Lewis Becker^73^, Rebecca Beer^85^, Amber Beitelshees^69^, Emelia Benjamin^86^, Takis Benos^87^, Marcos Bezerra^88^, Larry Bielak^64^, Thomas Blackwell^64^, Russell Bowler^89^, Ulrich Broeckel^90^, Jai Broome^70^, Deborah Brown^91^, Karen Bunting^63^, Esteban Burchard^92^, Carlos Bustamante^93^, Erin Buth^94^, Jonathan Cardwell^95^, Vincent Carey^96^, Julie Carrier^97^, Cara Carty^98^, Richard Casaburi^99^, Juan P Casas Romero^100^, James Casella^73^, Peter Castaldi^101^, Mark Chaffin^65^, Christy Chang^69^, Yi-Cheng Chang^102^, Daniel Chasman^103^, Sameer Chavan^95^, Bo-Juen Chen^63^, Wei-Min Chen^104^, Yii-Der Ida Chen^105^, Michael Cho^96^, Seung Hoan Choi^65^, Mina Chung^106^, Clary Clish^107^, Suzy Comhair^108^, Matthew Conomos^94^, Elaine Cornell^109^, Carolyn Crandall^99^, James Crapo^110^, L. Adrienne Cupples^111^, Jeffrey Curtis^64^, Brian Custer^112^, Coleen Damcott^69^, Dawood Darbar^113^, Sean David^114^, Colleen Davis^70^, Michelle Daya^95^, Mariza de Andrade^115^, Michael DeBaun^116^, Ranjan Deka^117^, Dawn DeMeo^96^, Scott Devine^69^, Huyen Dinh^78^, Harsha Doddapaneni^78^, Qing Duan^118^, Shannon Dugan-Perez^78^, Ravi Duggirala^119^, Jon Peter Durda^109^, Charles Eaton^120^, Lynette Ekunwe^71^, Adel El Boueiz^121^, Leslie Emery^70^, Serpil Erzurum^79^, Charles Farber^104^, Jesse Farek^78^, Tasha Fingerlin^122^, Matthew Flickinger^64^, Nora Franceschini^123^, Chris Frazar^70^, Mao Fu^69^, Stephanie M. Fullerton^70^, Lucinda Fulton^124^, Weiniu Gan^85^, Shanshan Gao^95^, Yan Gao^71^, Margery Gass^125^, Heather Geiger^126^, Bruce Gelb^127^, Mark Geraci^128^, Robert Gerszten^129^, Auyon Ghosh^96^, Chris Gignoux^75^, Mark Gladwin^87^, David Glahn^130^, Stephanie Gogarten^70^, Da-Wei Gong^69^, Harald Goring^131^, Sharon Graw^80^, Kathryn J. Gray^132^, Daniel Grine^95^, Colin Gross^64^, C. Charles Gu^124^, Yue Guan^69^, Namrata Gupta^65^, David M. Haas^133^, Jeff Haessler^125^, Michael Hall^134^, Yi Han^78^, Patrick Hanly^135^, Daniel Harris^136^, Nicola L. Hawley^137^, Ben Heavner^94^, Susan Heckbert^138^, Ryan Hernandez^92^, David Herrington^139^, Craig Hersh^140^, Bertha Hidalgo^141^, James Hixson^142^, Brian Hobbs^96^, John Hokanson^95^, Elliott Hong^69^, Karin Hoth^143^, Chao (Agnes) Hsiung^144^, Jianhong Hu^78^, Yi-Jen Hung^145^, Haley Huston^146^, Chii Min Hwu^147^, Rebecca Jackson^148^, Deepti Jain^70^, Cashell Jaquish^85^, Jill Johnsen^149^, Andrew Johnson^85^, Craig Johnson^70^, Rich Johnston^68^, Kimberly Jones^73^, Hyun Min Kang^150^, Shannon Kelly^151^, Eimear Kenny^127^, Michael Kessler^69^, Alyna Khan^70^, Ziad Khan^78^, Wonji Kim^152^, John Kimoff^153^, Greg Kinney^154^, Barbara Konkle^146^, Holly Kramer^155^, Christoph Lange^156^, Ethan Lange^95^, Cathy Laurie^70^, Cecelia Laurie^70^, Meryl LeBoff^96^, Jiwon Lee^96^, Sandra Lee^78^, Wen-Jane Lee^147^, Jonathon LeFaive^64^, David Levine^70^, Dan Levy^85^, Joshua Lewis^69^, Yun Li^118^, Henry Lin^105^, Honghuang Lin^157^, Simin Liu^158^, Yongmei Liu^159^, Yu Liu^160^, Kathryn Lunetta^157^, James Luo^85^, Ulysses Magalang^161^, Michael Mahaney^162^, Barry Make^73^, Alisa Manning^163^, JoAnn Manson^96^, Lisa Martin^164^, Melissa Marton^126^, Susan Mathai^95^, Susanne May^94^, Patrick McArdle^69^, Merry-Lynn McDonald^141^, Sean McFarland^152^, Daniel McGoldrick^165^, Caitlin McHugh^94^, Becky McNeil^166^, Hao Mei^71^, James Meigs^167^, Vipin Menon^78^, Luisa Mestroni^80^, Ginger Metcalf^78^, Deborah A Meyers^168^, Emmanuel Mignot^169^, Julie Mikulla^85^, Nancy Min^71^, Mollie Minear^170^, Ryan L Minster^87^, Matt Moll^101^, Zeineen Momin^78^, Courtney Montgomery^171^, Donna Muzny^78^, Josyf C Mychaleckyj^104^, Girish Nadkarni^127^, Rakhi Naik^73^, Sergei Nekhai^172^, Sarah C. Nelson^94^, Bonnie Neltner^95^, Caitlin Nessner^78^, Osuji Nkechinyere^78^, Jeff O’Connell^173^, Tim O’Connor^69^, Heather Ochs-Balcom^174^, Geoffrey Okwuonu^78^, Allan Pack^175^, David T. Paik^176^, James Pankow^177^, George Papanicolaou^85^, Cora Parker^178^, Juan Manuel Peralta^119^, Marco Perez^75^, James Perry^69^, Ulrike Peters^179^, Lawrence S Phillips^68^, Jacob Pleiness^64^, Toni Pollin^69^, Wendy Post^180^, Julia Powers Becker^181^, Meher Preethi Boorgula^95^, Michael Preuss^127^, Pankaj Qasba^85^, Dandi Qiao^96^, Zhaohui Qin^68^, Nicholas Rafaels^182^, Laura Raffield^183^, Mahitha Rajendran^78^, Vasan S. Ramachandran^157^, D.C. Rao^124^, Laura Rasmussen-Torvik^184^, Aakrosh Ratan^104^, Robert Reed^69^, Catherine Reeves^185^, Elizabeth Regan^110^, Alex Reiner^186^, Muagututi‘a Sefuiva Reupena^187^, Ken Rice^70^, Rebecca Robillard^188^, Nicolas Robine^126^, Dan Roden^189^, Carolina Roselli^65^, Ingo Ruczinski^73^, Alexi Runnels^126^, Pamela Russell^95^, Sarah Ruuska^146^, Kathleen Ryan^69^, Ester Cerdeira Sabino^190^, Danish Saleheen^191^, Shabnam Salimi^69^, Sejal Salvi^78^, Steven Salzberg^73^, Kevin Sandow^192^, Vijay G. Sankaran^193^, Jireh Santibanez^78^, Karen Schwander^124^, David Schwartz^95^, Frank Sciurba^87^, Christine Seidman^194^, Jonathan Seidman^195^, Frédéric Sériès^196^, Vivien Sheehan^197^, Stephanie L. Sherman^198^, Amol Shetty^69^, Aniket Shetty^95^, Wayne Hui-Heng Sheu^147^, M. Benjamin Shoemaker^199^, Brian Silver^200^, Edwin Silverman^96^, Robert Skomro^201^, Albert Vernon Smith^202^, Josh Smith^70^, Nicholas Smith^138^, Tanja Smith^63^, Sylvia Smoller^203^, Beverly Snively^204^, Michael Snyder^75^, Tamar Sofer^96^, Nona Sotoodehnia^70^, Adrienne M. Stilp^70^, Garrett Storm^205^, Elizabeth Streeten^69^, Jessica Lasky Su^96^, Yun Ju Sung^124^, Jody Sylvia^96^, Adam Szpiro^70^, Daniel Taliun^64^, Hua Tang^206^, Margaret Taub^73^, Matthew Taylor^80^, Simeon Taylor^69^, Marilyn Telen^74^, Timothy A. Thornton^70^, Machiko Threlkeld^207^, Lesley Tinker^125^, David Tirschwell^70^, Sarah Tishkoff^208^, Hemant Tiwari^209^, Catherine Tong^210^, Dhananjay Vaidya^73^, David Van Den Berg^211^, Peter VandeHaar^64^, Scott Vrieze^177^, Tarik Walker^95^, Robert Wallace^143^, Avram Walts^95^, Fei Fei Wang^70^, Heming Wang^212^, Jiongming Wang^202^, Karol Watson^99^, Jennifer Watt^78^, Daniel E. Weeks^87^, Joshua Weinstock^150^, Bruce Weir^70^, Scott T Weiss^213^, Lu-Chen Weng^214^, Jennifer Wessel^215^, Kayleen Williams^94^, L. Keoki Williams^216^, Carla Wilson^96^, James Wilson^217^, Lara Winterkorn^126^, Quenna Wong^70^, Joseph Wu^176^, Huichun Xu^69^, Ivana Yang^95^, Ketian Yu^64^, Seyedeh Maryam Zekavat^65^, Yingze Zhang^218^, Snow Xueyan Zhao^110^, Wei Zhao^219^, Xiaofeng Zhu^220^, Michael Zody^63^, Sebastian Zoellner^64^

63 - New York Genome Center, New York, New York, 10013; 64 - University of Michigan, Ann Arbor, Michigan, 48109; 65 - Broad Institute, Cambridge, Massachusetts, 02142; 66 - Cedars Sinai, Boston, Massachusetts, 02114; 67 - Children’s Hospital of Philadelphia, University of Pennsylvania, Philadelphia, Pennsylvania, 19104; 68 - Emory University, Atlanta, Georgia, 30322; 69 - University of Maryland, Baltimore, Maryland, 21201; 70 - University of Washington, Seattle, Washington, 98195; 71 - University of Mississippi, Jackson, Mississippi, 38677; 72 - National Institutes of Health, Bethesda, Maryland, 20892; 73 - Johns Hopkins University, Baltimore, Maryland, 21218; 74 - Duke University, Durham, North Carolina, 27708; 75 - Stanford University, Stanford, California, 94305; 76 - University of Wisconsin Milwaukee, Milwaukee, Wisconsin, 53211; 77 - Providence Health Care, Medicine, Vancouver; 78 - Baylor College of Medicine Human Genome Sequencing Center, Houston, Texas, 77030; 79 - Cleveland Clinic, Cleveland, Ohio, 44195; 80 - University of Colorado Anschutz Medical Campus, Aurora, Colorado, 80045; 81 - Columbia University, New York, New York, 10032; 82 - The Emmes Corporation, LTRC, Rockville, Maryland, 20850; 83 - Cleveland Clinic, Quantitative Health Sciences, Cleveland, Ohio, 44195; 84 - Johns Hopkins University, Medicine, Baltimore, Maryland, 21218; 85 - National Heart, Lung, and Blood Institute, National Institutes of Health, Bethesda, Maryland, 20892; 86 - Boston University, Massachusetts General Hospital, Boston University School of Medicine, Boston, Massachusetts, 02114; 87 - University of Pittsburgh, Pittsburgh, Pennsylvania, 15260; 88 - Fundação de Hematologia e Hemoterapia de Pernambuco - Hemope, Recife, 52011-000; 89 - National Jewish Health, National Jewish Health, Denver, Colorado, 80206; 90 - Medical College of Wisconsin, Milwaukee, Wisconsin, 53226; 91 - University of Texas Health at Houston, Pediatrics, Houston, Texas, 77030; 92 - University of California, San Francisco, San Francisco, California, 94143; 93 - Stanford University, Biomedical Data Science, Stanford, California, 94305; 94 - University of Washington, Biostatistics, Seattle, Washington, 98195; 95 - University of Colorado at Denver, Denver, Colorado, 80204; 96 - Brigham & Women’s Hospital, Boston, Massachusetts, 02115; 97 - University of Montreal; 98 - Washington State University, Pullman, Washington, 99164; 99 - University of California, Los Angeles, Los Angeles, California, 90095; 100 - Brigham & Women’s Hospital; 101 - Brigham & Women’s Hospital, Medicine, Boston, Massachusetts, 02115; 102 - National Taiwan University, Taipei, 10617; 103 - Brigham & Women’s Hospital, Division of Preventive Medicine, Boston, Massachusetts, 02215; 104 - University of Virginia, Charlottesville, Virginia, 22903; 105 - Lundquist Institute, Torrance, California, 90502; 106 - Cleveland Clinic, Cleveland Clinic, Cleveland, Ohio, 44195; 107 - Broad Institute, Metabolomics Platform, Cambridge, Massachusetts, 02142; 108 - Cleveland Clinic, Immunity and Immunology, Cleveland, Ohio, 44195; 109 - University of Vermont, Burlington, Vermont, 05405; 110 - National Jewish Health, Denver, Colorado, 80206; 111 - Boston University, Biostatistics, Boston, Massachusetts, 02115; 112 - Vitalant Research Institute, San Francisco, California, 94118; 113 - University of Illinois at Chicago, Chicago, Illinois, 60607; 114 - University of Chicago, Chicago, Illinois, 60637; 115 - Mayo Clinic, Health Quantitative Sciences Research, Rochester, Minnesota, 55905; 116 - Vanderbilt University, Nashville, Tennessee, 37235; 117 - University of Cincinnati, Cincinnati, Ohio, 45220; 118 - University of North Carolina, Chapel Hill, North Carolina, 27599; 119 - University of Texas Rio Grande Valley School of Medicine, Edinburg, Texas, 78539; 120 - Brown University, Providence, Rhode Island, 02912; 121 - Harvard University, Channing Division of Network Medicine, Cambridge, Massachusetts, 02138; 122 - National Jewish Health, Center for Genes, Environment and Health, Denver, Colorado, 80206; 123 - University of North Carolina, Epidemiology, Chapel Hill, North Carolina, 27599; 124 - Washington University in St Louis, St Louis, Missouri, 63130; 125 - Fred Hutchinson Cancer Research Center, Seattle, Washington, 98109; 126 - New York Genome Center, New York City, New York, 10013; 127 - Icahn School of Medicine at Mount Sinai, New York, New York, 10029; 128 - University of Pittsburgh, Pittsburgh, Pennsylvania; 129 - Beth Israel Deaconess Medical Center, Boston, Massachusetts, 02215; 130 - Boston Children’s Hospital, Harvard Medical School, Department of Psychiatry, Boston, Massachusetts, 02115; 131 - University of Texas Rio Grande Valley School of Medicine, San Antonio, Texas, 78229; 132 - Mass General Brigham, Obstetrics and Gynecology, Boston, Massachusetts, 02115; 133 - Indiana University, OB/GYN, Indianapolis, Indiana, 46202; 134 - University of Mississippi, Cardiology, Jackson, Mississippi, 39216; 135 - University of Calgary, Medicine, Calgary; 136 - University of Maryland, Genetics, Philadelphia, Pennsylvania, 19104; 137 - Yale University, Department of Chronic Disease Epidemiology, New Haven, Connecticut, 06520; 138 - University of Washington, Epidemiology, Seattle, Washington, 98195; 139 - Wake Forest Baptist Health, Winston-Salem, North Carolina, 27157; 140 - Brigham & Women’s Hospital, Channing Division of Network Medicine, Boston, Massachusetts, 02115; 141 - University of Alabama, Birmingham, Alabama, 35487; 142 - University of Texas Health at Houston, Houston, Texas, 77225; 143 - University of Iowa, Iowa City, Iowa, 52242; 144 - National Health Research Institute Taiwan, Institute of Population Health Sciences, NHRI, Miaoli County, 350; 145 - Tri-Service General Hospital National Defense Medical Center; 146 - Blood Works Northwest, Seattle, Washington, 98104; 147 - Taichung Veterans General Hospital Taiwan, Taichung City, 407; 148 - Oklahoma State University Medical Center, Internal Medicine, DIvision of Endocrinology, Diabetes and Metabolism, Columbus, Ohio, 43210; 149 - Blood Works Northwest, Research Institute, Seattle, Washington, 98104; 150 - University of Michigan, Biostatistics, Ann Arbor, Michigan, 48109; 151 - University of California, San Francisco, San Francisco, California, 94118; 152 - Harvard University, Cambridge, Massachusetts, 02138; 153 - McGill University, Montréal, QC H3A 0G4; 154 - University of Colorado at Denver, Epidemiology, Aurora, Colorado, 80045; 155 - Loyola University, Public Health Sciences, Maywood, Illinois, 60153; 156 - Harvard School of Public Health, Biostats, Boston, Massachusetts, 02115; 157 - Boston University, Boston, Massachusetts, 02215; 158 - Brown University, Epidemiology and Medicine, Providence, Rhode Island, 02912; 159 - Duke University, Cardiology, Durham, North Carolina, 27708; 160 - Stanford University, Cardiovascular Institute, Stanford, California, 94305; 161 - Ohio State University, Division of Pulmonary, Critical Care and Sleep Medicine, Columbus, Ohio, 43210; 162 - University of Texas Rio Grande Valley School of Medicine, Brownsville, Texas, 78520; 163 - Broad Institute, Harvard University, Massachusetts General Hospital; 164 - George Washington University, cardiology, Washington, District of Columbia, 20037; 165 - University of Washington, Genome Sciences, Seattle, Washington, 98195; 166 - RTI International; 167 - Massachusetts General Hospital, Medicine, Boston, Massachusetts, 02114; 168 - University of Arizona, Tucson, Arizona, 85721; 169 - Stanford University, Center For Sleep Sciences and Medicine, Palo Alto, California, 94304; 170 - National Institute of Child Health and Human Development, National Institutes of Health, Bethesda, Maryland, 20892; 171 - Oklahoma Medical Research Foundation, Genes and Human Disease, Oklahoma City, Oklahoma, 73104; 172 - Howard University, Washington, District of Columbia, 20059; 173 - University of Maryland, Balitmore, Maryland, 21201; 174 - University at Buffalo, Buffalo, New York, 14260; 175 - University of Pennsylvania, Division of Sleep Medicine/Department of Medicine, Philadelphia, Pennsylvania, 19104-3403; 176 - Stanford University, Stanford Cardiovascular Institute, Stanford, California, 94305; 177 - University of Minnesota, Minneapolis, Minnesota, 55455; 178 - RTI International, Biostatistics and Epidemiology Division, Research Triangle Park, North Carolina, 27709- 2194; 179 - Fred Hutchinson Cancer Research Center, Fred Hutch and UW, Seattle, Washington, 98109; 180 - Johns Hopkins University, Cardiology/Medicine, Baltimore, Maryland, 21218; 181 - University of Colorado at Denver, Medicine, Denver, Colorado, 80204; 182 - University of Colorado at Denver, Denver, Colorado, 80045; 183 - University of North Carolina, Genetics, Chapel Hill, North Carolina, 27599; 184 - Northwestern University, Chicago, Illinois, 60208; 185 - New York Genome Center, New York Genome Center, New York City, New York, 10013; 186 - Fred Hutchinson Cancer Research Center, University of Washington, Seattle, Washington, 98109; 187 - Lutia I Puava Ae Mapu I Fagalele, Apia; 188 - University of Ottawa, Sleep Research Unit, University of Ottawa Institute for Mental Health Research, Ottawa, ON K1Z 7K4; 189 - Vanderbilt University, Medicine, Pharmacology, Biomedicla Informatics, Nashville, Tennessee, 37235; 190 - Universidade de Sao Paulo, Faculdade de Medicina, Sao Paulo, 01310000; 191 - Columbia University, New York, New York, 10027; 192 - Lundquist Institute, TGPS, Torrance, California, 90502; 193 - Harvard University, Division of Hematology/Oncology, Boston, Massachusetts, 02115; 194 - Harvard Medical School, Genetics, Boston, Massachusetts, 02115; 195 - Harvard Medical School, Boston, Massachusetts, 02115; 196 - Université Laval, Quebec City, G1V 0A6; 197 - Emory University, Pediatrics, Atlanta, Georgia, 30307; 198 - Emory University, Human Genetics, Atlanta, Georgia, 30322; 199 - Vanderbilt University, Medicine/Cardiology, Nashville, Tennessee, 37235; 200 - UMass Memorial Medical Center, Worcester, Massachusetts, 01655; 201 - University of Saskatchewan, Saskatoon, SK S7N 5C9; 202 - University of Michigan; 203 - Albert Einstein College of Medicine, New York, New York, 10461; 204 - Wake Forest Baptist Health, Biostatistical Sciences, Winston-Salem, North Carolina, 27157; 205 - University of Colorado at Denver, Genomic Cardiology, Aurora, Colorado, 80045; 206 - Stanford University, Genetics, Stanford, California, 94305; 207 - University of Washington, University of Washington, Department of Genome Sciences, Seattle, Washington, 98195; 208 - University of Pennsylvania, Genetics, Philadelphia, Pennsylvania, 19104; 209 - University of Alabama, Biostatistics, Birmingham, Alabama, 35487; 210 - University of Washington, Department of Biostatistics, Seattle, Washington, 98195; 211 - University of Southern California, USC Methylation Characterization Center, University of Southern California, California, 90033; 212 - Brigham & Women’s Hospital, Mass General Brigham, Boston, Massachusetts, 02115; 213 - Brigham & Women’s Hospital, Channing Division of Network Medicine, Department of Medicine, Boston, Massachusetts, 02115; 214 - Massachusetts General Hospital, Boston, Massachusetts, 02114; 215 - Indiana University, Epidemiology, Indianapolis, Indiana, 46202; 216 - Henry Ford Health System, Detroit, Michigan, 48202; 217 - Beth Israel Deaconess Medical Center, Cardiology, Cambridge, Massachusetts, 02139; 218 - University of Pittsburgh, Medicine, Pittsburgh, Pennsylvania, 15260; 219 - University of Michigan, Department of Epidemiology, Ann Arbor, Michigan, 48109; 220 - Case Western Reserve University, Department of Population and Quantitative Health Sciences, Cleveland, Ohio, 44106

