## Supplementary Text for "Whole genome sequence analysis of blood lipid levels in >66,000 individuals"

| **Table of Contents** |  |
| --- | --- |
| **Supplementary Text: Cohort description** | **Page No** |
| Old Order Amish (Amish, n=1,083) | 3 |
| Atherosclerosis Risk in Communities study (ARIC, n=8,016) | 3 |
| Mt Sinai BioMe Biobank (BioMe, n=9,848) | 3 |
| Coronary Artery Risk Development in Young Adults (CARDIA, n=3,056) | 4 |
| Cleveland Family Study (CFS, n=579) | 4 |
| Cardiovascular Health Study (CHS, n=3,456) | 5 |
| Diabetes Heart Study (DHS, n=365) | 5 |
| Framingham Heart Study (FHS, n=3,992) | 6 |
| Genetic Studies of Atherosclerosis Risk (GeneSTAR, n=1,757) | 6 |
| Genetic Epidemiology Network of Arteriopathy (GENOA, n=1,046) | 7 |
| Genetic Epidemiology Network of Salt Sensitivity (GenSalt, n=1,772) | 7 |
| Genetics of Lipid-Lowering Drugs and Diet Network (GOLDN, n=926) | 7 |
| Hispanic Community Health Study - Study of Latinos (HCHS_SOL, n=7714) | 8 |
| Hypertension Genetic Epidemiology Network and Genetic Epidemiology Network of Arteriopathy (HyperGEN, n=1,853) | 8 |
| Jackson Heart Study (JHS, n=2,847) | 9 |
| Multi-Ethnic Study of Atherosclerosis (MESA, n=5,290) | 9 |
| Massachusetts General Hospital Atrial Fibrillation Study (MGH_AF, n=683) | 10 |
| San Antonio Family Study (SAFS, n=619) | 10 |
| Samoan Adiposity Study (SAS, n=1,182) | 11 |
| Taiwan Study of Hypertension using Rare Variants (THRV, n=1,982) | 11 |
| Women’s Health Initiative (WHI, n=8,263) | 12 |
| **TOPMed Omics Support table** | 13 |
| **Supplementary Figures: Legends** |  |
| Supplementary Fig. 1: Overview of the workflow implement in the study | 16 |
| Supplementary Fig. 2: Manhattan plots for the four lipid phenotypes | 16 |
| Supplementary Fig. 3: Correlation of CETP gene expression effects with HDL-C effects | 16 |
| Supplementary Fig. 4 Genome wide view of coding and non-coding aggregate sets | 16 |
| Supplementary Fig. 5 Conditional analysis of coding rare-variants from the same gene and a near-by gene | 17 |
| Supplementary Fig. 6 Distribution of Mendelian gene STAAR-O p-values | 17 |
| **Supplementary Tables: Legends** |  |
| Supplementary Table 1: TOPMed freeze 8 phenotype data distributed based on 21 cohorts | 18 |
| Supplementary Table 2: TOPMed Freeze 8 genotype distribution based on chromosome | 18 |
| Supplementary Table 3: TOPMed Freeze 8 variants summary | 18 |
| Supplementary Table 4: Ancestry specificity and replication of putative novel variants | 18 |
| Supplementary Table 5: Baseline characteristics of replication cohorts | 19 |
| Supplementary Table 6: Evaluation of suggestive lipid alleles in TOPMed with independent datasets | 19 |
| Supplementary Table 7: List of CETP variants significant at suggestive p-value (5X10-07) from both LDL-C and HDL-C GWAS | 19 |
| Supplementary Table 8: Significant gene-centric coding rare variant aggregate sets | 19 |
| Supplementary Table 9: Significant gene-centric non-coding rare variant aggregate sets | 19 |
| Supplementary Table 10: Significant gene-centric coding rare variant aggregate sets after conditional analysis | 20 |
| Supplementary Table 11: Significant gene-centric non-coding rare variant aggregate sets after conditional analysis | 20 |
| Supplementary Table 12: Region-based sliding-window results before and after conditional analysis | 20 |
| Supplementary Table 13: Region-based dynamic-window results before and after conditional analysis | 20 |
| Supplementary Table 14: Significant rare non-coding variants after conditioning on rare coding variants | 21 |
| Supplementary Table 15: Common variants and rare variant aggregates at Mendelian lipid genes | 21 |
| **Supplementary Figures** | 22-35 |
| **References** | 36-37 |

**Supplementary Text:**

**Old Order Amish (Amish, n=1,083):**

*TOPMed dbGaP accession#: phs000956, Parent dbGaP accession#: phs000391, Sequencing Center: Broad Institute of MIT and Harvard.*

The Old Order Amish (OOA) population of Lancaster County, PA immigrated to the Colonies from Western Europe in the early 1700’s. Investigators at University of Maryland School of Medicine have been studying the genetic determinants of cardiometabolic health in this population since 1993. To date, over 7,000 Amish adults have participated in one or more of our studies. The Heredity and Phenotype Intervention (HAPI) Heart Study was initiated in 2002 and chose to study the OOA to test the genetic effects of complex phenotypes from this genetically homogeneous population^1^. The Amish Research Group includes investigators with a diverse range of interests in population and basic science and in clinical and translational research. Whole genome sequencing (WGS) for the Trans-Omics in Precision Medicine (TOPMed) program was supported by the National Heart, Lung and Blood Institute (NHLBI). The Amish studies were supported by NIH grants R01 AG18728, U01 HL072515, R01 HL088119, R01 HL121007, and P30 DK072488.

**Atherosclerosis Risk in Communities study (ARIC, n=8,016):**

*TOPMed dbGaP accession#: phs001211, Parent dbGaP accession#: phs000280, Sequencing Center: Baylor College of Medicine Human Genome Sequencing Center and Broad Institute of MIT and Harvard.*

The ARIC study is a population-based prospective cohort study of cardiovascular disease sponsored by the National Heart, Lung, and Blood Institute (NHLBI). ARIC included 15,792 individuals, predominantly European American and African American, aged 45-64 years at baseline (1987-89), chosen by probability sampling from four US communities. Cohort members completed three additional triennial follow-up examinations, a fifth exam in 2011-2013, a sixth exam in 2016-2017, a seventh exam in 2018-2019, and an eight exam in 2020. The ARIC study has been described in detail previously^2^.

Whole genome sequencing (WGS) for the Trans-Omics in Precision Medicine (TOPMed) program was supported by the National Heart, Lung and Blood Institute (NHLBI). WGS for “NHLBI TOPMed: Atherosclerosis Risk in Communities (ARIC)” (phs001211) was performed at the Baylor College of Medicine Human Genome Sequencing Center (HHSN268201500015C and 3U54HG003273-12S2) and the Broad Institute for MIT and Harvard (3R01HL092577-06S1). Centralized read mapping and genotype calling, along with variant quality metrics and filtering were provided by the TOPMed Informatics Research Center (3R01HL-117626-02S1). Phenotype harmonization, data management, sample-identity QC, and general study coordination, were provided by the TOPMed Data Coordinating Center (3R01HL-120393-02S1). We gratefully acknowledge the studies and participants who provided biological samples and data for TOPMed.

The Genome Sequencing Program (GSP) was funded by the National Human Genome Research Institute (NHGRI), the National Heart, Lung, and Blood Institute (NHLBI), and the National Eye Institute (NEI). The GSP Coordinating Center (U24 HG008956) contributed to cross-program scientific initiatives and provided logistical and general study coordination. The Centers for Common Disease Genomics (CCDG) program was supported by NHGRI and NHLBI, and whole genome sequencing was performed at the Baylor College of Medicine Human Genome Sequencing Center (UM1 HG008898 and R01HL059367).

The Atherosclerosis Risk in Communities study has been funded in whole or in part with Federal funds from the National Heart, Lung, and Blood Institute, National Institutes of Health, Department of Health and Human Services (contract numbers HHSN268201700001I, HHSN268201700002I, HHSN268201700003I, HHSN268201700004I and HHSN268201700005I). The authors thank the staff and participants of the ARIC study for their important contributions.

**Mt Sinai BioMe Biobank (BioMe, n=9,848):**

*TOPMed dbGaP accession#: phs001644, Parent dbGaP accession#: phs000925, Sequencing Center: Baylor College of Medicine Human Genome Sequencing Center, Northwest Genomics Center*

The Institute for Personalized Medicine at the Icahn School of Medicine at Mount Sinai is leading the movement toward diagnosis and classification of disease according to the patient’s molecular profile. BioMe is the major effort towards this goal, BioMe, an electronic medical record-linked biobank that enables researchers to conduct genetic, epidemiologic, molecular, and genomic studies rapidly and efficiently on large collections of research specimens linked with medical information. BioMed cohort is composed of White, Black, Hispanic and Asian populations. The Mount Sinai Medical Center services diverse local communities of upper Manhattan, including Central Harlem (86% African American), East Harlem (88% Hispanic Latino), and Upper East Side (88% Caucasian/white) with broad health disparities^3^. Whole genome sequencing (WGS) for the Trans-Omics in Precision Medicine (TOPMed) program was supported by the National Heart, Lung and Blood Institute (NHLBI). The Mount Sinai BioMe Biobank has been supported by The Andrea and Charles Bronfman Philanthropies and in part by Federal funds from the NHLBI and NHGRI (U01HG00638001; U01HG007417; X01HL134588).

**Coronary Artery Risk Development in Young Adults (CARDIA, n=3,056):**

TOPMed dbGaP accession#: phs001612, Parent dbGaP accession#: phs000285, *Sequencing Center: Baylor College of Medicine Human Genome Sequencing Center.*

The Coronary Artery Risk Development in Young Adults (CARDIA) Study is a study examining the development and determinants of clinical and subclinical cardiovascular disease and their risk factors. It began in 1985-6 with a group of 5115 black and white men and women aged 18-30 years^4^. The participants were selected so that there would be approximately the same number of people in subgroups of race, gender, education (high school or less and more than high school) and age (18-24 and 25-30) in each of 4 centers: Birmingham, AL; Chicago, IL; Minneapolis, MN; and Oakland, CA. These same participants were asked to participate in follow-up examinations during 1987-1988 (Year 2), 1990-1991 (Year 5), 1992-1993 (Year 7), 1995-1996 (Year 10), 2000-2001 (Year 15), 2005-2006 (Year 20), 2010-2011 (Year 25), and 2015-2016 (Year 30). CARDIA cohort is composed of White and Black populations. Whole genome sequencing (WGS) for the Trans-Omics in Precision Medicine (TOPMed) program was supported by the National Heart, Lung and Blood Institute (NHLBI). The Coronary Artery Risk Development in Young Adults Study (CARDIA) is conducted and supported by the National Heart, Lung, and Blood Institute (NHLBI) in collaboration with the University of Alabama at Birmingham (HHSN268201800005I & HHSN268201800007I), Northwestern University (HHSN268201800003I), University of Minnesota (HHSN268201800006I), and Kaiser Foundation Research Institute (HHSN268201800004I). CARDIA was also partially supported by the Intramural Research Program of the National Institute on Aging (NIA) and an intra‐agency agreement between NIA and NHLBI (AG0005).

**Cleveland Family Study (CFS, n=579):**

*TOPMed dbGaP accession#: phs000954, Parent dbGaP accession#: phs000284, Sequencing Center:* *McDonnell Genome Institute (MGI) at Washington University.*

The CFS is a family-based longitudinal study that includes participants with laboratory diagnosed sleep apnea, their family members and neighborhood control families followed between 1990 and 2006^5^. After an overnight fast, blood was collected which was assayed for lipid levels at the University of Vermont Laboratory for Clinical Biochemistry Research. Lipids (triglycerides, HDL cholesterol) from fasted blood serum were measured by enzymatic methods. CFS cohort is composed of White and Black populations. Whole genome sequencing (WGS) for the Trans-Omics in Precision Medicine (TOPMed) program was supported by the National Heart, Lung and Blood Institute (NHLBI). CFS is supported by grants from the NHLBI (HL046389, HL113338, and 1R35HL135818).

**Cardiovascular Health Study (CHS, n=3,456):**

*TOPMed dbGaP accession#: phs001368, Parent dbGaP accession#: phs000287, Sequencing Center: Baylor College of Medicine Human Genome Sequencing Center.*

The Cardiovascular Health Study (CHS) is an NHLBI-funded observational study of risk factors for cardiovascular disease in adults 65 years or older. Starting in 1989, and continuing through 1999, participants underwent annual extensive clinical examinations. Measurements included traditional risk factors such as blood pressure and lipids. Additionally, measures of subclinical disease, including echocardiography of the heart, carotid ultrasound, and cranial magnetic-resonance imaging (MRI) were determined. At six-month intervals between clinic visits, and once clinic visits ended, participants were contacted by phone to ascertain hospitalizations and health status. The main outcomes are coronary heart disease (CHD), angina, heart failure (HF), stroke, transient ischemic attack (TIA), claudication, and mortality^6^. Participants continue to be contacted by phone every 6 months. Participants from four counties were included in the study: Forsyth County, North Carolina; Sacramento County, California; Washington County, Maryland; and Pittsburgh, Pennsylvania. CHS cohort is composed of White and Black populations. Whole genome sequencing (WGS) for the Trans-Omics in Precision Medicine (TOPMed) program was supported by the National Heart, Lung and Blood Institute (NHLBI). The CHS research was supported by NHLBI contracts 75N92021D00006, HHSN268201200036C, HHSN268200800007C, HHSN268201800001C, N01HC55222, N01HC85079, N01HC85080, N01HC85081, N01HC85082, N01HC85083, N01HC85086; and NHLBI grants U01HL080295, R01HL087652, R01HL105756, R01HL103612, R01HL120393, R01HL130114, and R01 HL059367, with additional contribution from the National Institute of Neurological Disorders and Stroke (NINDS). Additional support was provided through R01AG023629 from the National Institute on Aging (NIA).

**Diabetes Heart Study (DHS, n=365):**

*TOPMed dbGaP accession#: phs001412, Parent dbGaP accession#: phs001012, Sequencing Center: Broad Institute of MIT and Harvard*

The Diabetes Heart Study (DHS) is a family-based study enriched for type 2 diabetes (T2D). The cohort was recruited between 1998 and 2006. Participants were extensively phenotyped for measures of subclinical CVD and other known CVD risk factors^7^. Primary outcomes were quantified burden of vascular calcified plaque in the coronary artery, carotid artery, and abdominal aorta all determined from non-contrast computed tomography scans. Whole genome sequencing (WGS) for the Trans-Omics in Precision Medicine (TOPMed) program was supported by the National Heart, Lung and Blood Institute (NHLBI). The DHS research was supported by R01 HL92301, R01 HL67348, R01 NS058700, R01 AR48797, R01 DK071891, R01 AG058921, the General Clinical Research Center of the Wake Forest University School of Medicine (M01 RR07122, F32 HL085989), the American Diabetes Association, and a pilot grant from the Claude Pepper Older Americans Independence Center of Wake Forest University Health Sciences (P60 AG10484).

**Framingham Heart Study (FHS, n=3,992):**

*TOPMed dbGaP accession#: phs000974, Parent dbGaP accession#: phs000007, Sequencing Center: Broad Institute of MIT and Harvard*

The Framingham Heart Study (FHS) is a prospective cohort study of 3 generations of subjects who have been followed up to 65 years to evaluate risk factors for cardiovascular disease.^13-16^ Its large sample of ~15,000 men and women who have been extensively phenotyped with repeated examinations make it ideal for the study of genetic associations with cardiovascular disease risk factors and outcomes. DNA samples have been collected and immortalized since the mid-1990s and are available on ~8000 study participants in 1037 families. These samples have been used for collection of GWAS array data and exome chip data in nearly all with DNA samples, and for targeted sequencing, deep exome sequencing and light coverage whole genome sequencing in limited numbers. Additionally, mRNA and miRNA expression data, DNA methylation data, metabolomics and other 'omics data are available on a sizable portion of study participants. This project on the focuses on deep whole genome sequencing (mean 30X coverage) in 3,992 subjects with lipid data available.

FHS acknowledges the support of contracts NO1-HC-25195 and HHSN268201500001I from the National Heart, Lung and Blood Institute and grant supplement R01 HL092577-06S1 for this research. WGS for “NHLBI TOPMed: Whole Genome Sequencing and Related Phenotypes in the Framingham Heart Study” (phs000974) was performed at the Broad Institute of MIT and Harvard (HHSN268201500014C, 3R01HL092577-06S1, and 3U54HG003067-12S2). We also acknowledge the dedication of the FHS study participants without whom this research would not be possible.

**Genetic Studies of Atherosclerosis Risk (GeneSTAR, n=1,757):**

*TOPMed dbGaP accession#: phs001218, Parent dbGaP accession#: phs000375, Sequencing Center: Broad Institute of MIT and Harvard, Illumina Genomic Services, PSOMAGEN (formerly Macrogen).*

GeneSTAR is a family-based study in initially healthy brothers and sisters, and offspring of people with early-onset coronary disease. The goal is to discover and amplify mechanisms of stroke and coronary heart disease. In 1982 GeneSTAR was created to study patterns of coronary heart disease and related risk factors in families with early-onset coronary disease, identified from10 Baltimore area Hospitals. Extensive additional cardiovascular testing and risk assessment was done at baseline and serially^12,13^. Follow-up was carried out to determine incident cardiovascular disease, stroke, peripheral arterial disease, diabetes, cancer, and related comorbidities, from 5 to 30 years after study entry. Whole genome sequencing (WGS) for the Trans-Omics in Precision Medicine (TOPMed) program was supported by the National Heart, Lung and Blood Institute (NHLBI). GeneSTAR was supported by grants from the National Institutes of Health/National Heart, Lung, and Blood Institute (U01 HL72518, HL087698, HL49762, HL58625, HL071025, HL112064), the National Institutes of Health/National Institute of Nursing Research (NR0224103), and by a grant from the National Institutes of Health/National Center for Research Resources (M01-RR000052) to the Johns Hopkins General Clinical Research Center.

**Genetic Epidemiology Network of Arteriopathy (GENOA, n=1,046):**

*TOPMed dbGaP accession#: phs001345, Parent dbGaP accession#: phs001238, Sequencing Center: Broad Institute of MIT and Harvard, McDonnell Genome Institute (MGI) at Washington University.*

The Genetic Epidemiology Network of Arteriopathy (GENOA) is one of four networks in the NHLBI Family-Blood Pressure Program (FBPP)^14^. GENOA's long-term objective is to elucidate the genetics of target organ complications of hypertension, including both atherosclerotic and arteriolosclerotic complications involving the heart, brain, kidneys, and peripheral arteries^15^. The longitudinal GENOA Study recruited European-American and African-American sibships with at least 2 individuals with clinically diagnosed essential hypertension before age 60 years. All other members of the sibship were invited to participate regardless of their hypertension status. Participants were diagnosed with hypertension if they had either 1) a previous clinical diagnosis of hypertension by a physician with current anti-hypertensive treatment, or 2) an average systolic blood pressure ≥ 140 mm Hg or diastolic blood pressure ≥ 90 mm Hg based on the second and third readings at the time of their clinic visit. Only participants of the African-American Cohort were sequenced through TOPMed. Whole genome sequencing (WGS) for the Trans-Omics in Precision Medicine (TOPMed) program was supported by the National Heart, Lung and Blood Institute (NHLBI). Support for GENOA was provided by the National Heart, Lung and Blood Institute (HL054457, HL054464, HL054481, and HL087660) of the National Institutes of Health.

**Genetic Epidemiology Network of Salt Sensitivity (GenSalt, n=1,772):**

*TOPMed dbGaP accession#: phs001217, Parent dbGaP accession#: phs000784, Sequencing Center: Broad Institute of MIT and Harvard.*

GenSalt utilizes a family feeding-study design. Each family is ascertained through a proband with untreated prehypertension or stage-1 hypertension in rural China. Medical history, lifestyle risk factors, and cold pressor tests are obtained at baseline visits while BP, weight, blood and urine specimens are collected at baseline and follow-up visits^16^. The dietary intervention includes a 7-day low sodium-feeding (51.3 mmol/day), a 7-day high sodium-feeding (307.8 mmol/day), and a 7-day high sodium-feeding with an oral potassium supplementation (60 mmol/day). Whole genome sequencing (WGS) for the Trans-Omics in Precision Medicine (TOPMed) program was supported by the National Heart, Lung and Blood Institute (NHLBI). GenSalt was supported by research grants (U01HL072507, R01HL087263, and R01HL090682) from the National Heart, Lung and Blood Institute, National Institutes of Health, Bethesda, MD.

**Genetics of Lipid-Lowering Drugs and Diet Network (GOLDN, n=926):**

*TOPMed dbGaP accession#: phs001359, Parent dbGaP accession#: phs000741, Sequencing Center: McDonnell Genome Institute (MGI) at Washington University.*

GOLDN is a family-based study of European descent individuals recruited in Minneapolis and Salt Lake City (two of the NHLBI Family Heart Study sites). Contributions of genes, shared and individual environments, and behaviors to variations in risk factors, preclinical atherosclerosis, and CHD were estimated in the study^17^. It aims to uncover genetic predictors of variability in lipid phenotypes, which include both fasting and postprandial lipids quantified using traditional methods, NMR, and high throughput lipidomics. Whole genome sequencing (WGS) for the Trans-Omics in Precision Medicine (TOPMed) program was supported by the National Heart, Lung and Blood Institute (NHLBI). GOLDN biospecimens, baseline phenotype data, and intervention phenotype data were collected with funding from National Heart, Lung and Blood Institute (NHLBI) grant U01 HL072524.

**Hispanic Community Health Study - Study of Latinos (HCHS_SOL, n=7714):**

*TOPMed dbGaP accession#: phs001395, Parent dbGaP accession#: phs000810, Sequencing Center: Baylor College of Medicine Human Genome Sequencing Center.*

The Hispanic Community Health Study (HCHS)/Study of Latinos (SOL) is a multicenter, community-based cohort study of Hispanic/Latino adults in the United States. The main goal of the study is to identify risk factors which could either be protective or harmful to the Hispanic community^18,19^. A total of 16,000 Hispanic/Latino individuals of age 18-74 were recruited. Participants are recruited in community areas surrounding four field centers in the Bronx, Chicago, Miami, and San Diego. Whole genome sequencing (WGS) for the Trans-Omics in Precision Medicine (TOPMed) program was supported by the National Heart, Lung and Blood Institute (NHLBI). The Hispanic Community Health Study/Study of Latinos was carried out as a collaborative study supported by contracts from the National Heart, Lung, and Blood Institute (NHLBI) to the University of North Carolina (N01-HC65233), University of Miami (N01-HC65234), Albert Einstein College of Medicine (N01-HC65235), Northwestern University (N01-HC65236), and San Diego State University (N01-HC65237).

**Hypertension Genetic Epidemiology Network and Genetic Epidemiology Network of Arteriopathy (HyperGEN, n=1,853):**

*TOPMed dbGaP accession#: phs001293, Parent dbGaP accession#: phs001293, Sequencing Center: McDonnell Genome Institute (MGI) at Washington University.*

The Hypertension Genetic Epidemiology Network Study (HyperGEN) - Genetics of Left Ventricular (LV) Hypertrophy is a familial study aimed to understand genetic risk factors for LV hypertrophy by conducting genetic studies of continuous traits from echocardiography exams ^20,21^. As part of HyperGEN study, four field centers recruited African American and white hypertensive siblings, aged 23 to 87 years. Data from detailed clinical exams as well as genotyping data for linkage studies, candidate gene studies and GWAS have been collected and is shared between HyperGEN and the ancillary HyperGEN - Genetics of LV Hypertrophy study. Whole genome sequencing (WGS) for the Trans-Omics in Precision Medicine (TOPMed) program was supported by the National Heart, Lung and Blood Institute (NHLBI). The HyperGEN Study is part of the National Heart, Lung, and Blood Institute (NHLBI) Family Blood Pressure Program; collection of the data represented here was supported by grants U01 HL054472 (MN Lab), U01 HL054473 (DCC), U01 HL054495 (AL FC), and U01 HL054509 (NC FC). The HyperGEN: Genetics of Left Ventricular Hypertrophy Study was supported by NHLBI grant R01 HL055673 with whole-genome sequencing made possible by supplement -18S1.

**Jackson Heart Study (JHS, n=2,847):**

*TOPMed dbGaP accession#: phs000964, Parent dbGaP accession#: phs000286, Sequencing Center: McDonnell Genome Institute (MGI) at Washington University.*

The JHS is a large, community-based, observational study, four subsamples of participants (random, volunteer, ARIC (continuing from Atherosclerosis Risk in Communities study), and family) residing in the Jackson, Mississippi metropolitan statistical area (MSA) were included^22,23,24^. Recruitment was limited to persons 35-84 years old except in the family cohort, where those 21 years old and above were eligible. Participants provided extensive medical and social history, had an array of physical and biochemical measurements and diagnostic procedures, and provided genomic DNA. Whole genome sequencing (WGS) for the Trans-Omics in Precision Medicine (TOPMed) program was supported by the National Heart, Lung and Blood Institute (NHLBI). The Jackson Heart Study (JHS) is supported and conducted in collaboration with Jackson State University (HHSN268201800013I), Tougaloo College (HHSN268201800014I), the Mississippi State Department of Health (HHSN268201800015I) and the University of Mississippi Medical Center (HHSN268201800010I, HHSN268201800011I and HHSN268201800012I) contracts from the National Heart, Lung, and Blood Institute (NHLBI) and the National Institute on Minority Health and Health Disparities (NIMHD).

**Multi-Ethnic Study of Atherosclerosis (MESA, n=5,290):**

*TOPMed dbGaP accession#: phs001416, Parent dbGaP accession#: phs000209,*

*Sequencing Center: Broad Institute of MIT and Harvard.*

The Multi-Ethnic Study of Atherosclerosis (MESA) is a study of the characteristics of subclinical cardiovascular disease (disease detected non-invasively before it has produced clinical signs and symptoms) and the risk factors that predict progression to clinically overt cardiovascular disease or progression of the subclinical disease. MESA researchers study a diverse, population-based sample of 6,814 asymptomatic men and women aged 45-84 from six field centers across the United States. Approximately 38 % of the recruited participants are white, 28 % African-American, 22 % Hispanic, and 12 % Asian, predominantly of Chinese descent^25^. Six exams have been completed since 2000. Participants are contacted every 9 to 12 months throughout the study to assess clinical morbidity and mortality. The final 18 months of the study will be dedicated to close out and data analysis and publication. Baseline measurements will include measurement of coronary calcium using computed tomography; measurement of ventricular mass and function using cardiac magnetic resonance imaging; measurement of flow-mediated brachial artery endothelial vasodilation, carotid intimal-medial wall thickness, and distensibility of the carotid arteries using ultrasonography; measurement of peripheral vascular disease using ankle and brachial blood pressures; electrocardiography; and assessments of microalbuminuria, standard CVD risk factors, sociodemographic factors, life habits, and psychosocial factors.

Centralized read mapping and genotype calling, along with variant quality metrics and filtering were provided by the TOPMed Informatics Research Center (3R01HL-117626-02S1). Phenotype harmonization, data management, sample-identity QC, and general study coordination, were provided by the TOPMed Data Coordinating Center (3R01HL-120393-02S1). MESA and the MESA SHARe project are conducted and supported by the National Heart, Lung, and Blood Institute (NHLBI) in collaboration with MESA investigators. Whole genome sequencing (WGS) for the Trans-Omics in Precision Medicine (TOPMed) program was supported by the National Heart, Lung and Blood Institute (NHLBI). Support for MESA is provided by contracts HHSN268201500003I, N01-HC-95159, N01-HC-95160, N01-HC-95161, N01-HC-95162, N01-HC-95163, N01-HC-95164, N01-HC-95165, N01-HC-95166, N01-HC-95167, N01-HC-95168, N01-HC-95169, UL1-TR-000040, UL1-TR-001079, UL1-TR-001420. MESA Family is conducted and supported by the National Heart, Lung, and Blood Institute (NHLBI) in collaboration with MESA investigators. Support is provided by grants and contracts R01HL071051, R01HL071205, R01HL071250, R01HL071251, R01HL071258, R01HL071259, and by the National Center for Research Resources, Grant UL1RR033176. The provision of genotyping data was supported in part by the National Center for Advancing Translational Sciences, CTSI grant UL1TR001881, and the National Institute of Diabetes and Digestive and Kidney Disease Diabetes Research Center (DRC) grant DK063491 to the Southern California Diabetes Endocrinology Research Center.

**Massachusetts General Hospital Atrial Fibrillation Study (MGH_AF, n=683):**

*TOPMed dbGaP accession#: phs001062, Parent dbGaP accession#: phs001001*

*Sequencing Center: Broad Institute of MIT and Harvard.*

The MGH-AF study was initiated to study the effects of atrial fibrillations by comparing unaffected and affected family members^26,27^. The participants provide details on past medical history, AF treatment and family history. An electrocardiogram is performed; the results of an echocardiogram are obtained along with blood samples. For the TOPMed whole genome sequencing project only early-onset atrial fibrillation cases were sequenced. Early-onset atrial fibrillation was defined as an age of onset prior to 66 years of age. Whole genome sequencing (WGS) for the Trans-Omics in Precision Medicine (TOPMed) program was supported by the National Heart, Lung and Blood Institute (NHLBI). The MGH AF Study was supported by grants to Dr. Ellinor from the Fondation Leducq (14CVD01), the National Institutes of Health to Dr. Ellinor (1RO1HL092577, R01HL128914, K24HL105780) and Dr. Lubitz (1R01HL139731) and by grants from the American Heart Association to Dr. Ellinor (18SFRN34110082) and to Dr. Lubitz (18SFRN34250007).

**San Antonio Family Study (SAFS, n=619):**

*TOPMed dbGaP accession#: phs001215, Parent dbGaP accession#: phs000462, Sequencing Center: Illumina Genomic Services.*

The SAFS began in 1991, and included 1,431 individuals in 42 extended families at baseline. Probands were 40- to 60-year-old low-income Mexican Americans selected at random without regard to presence or absence of disease, almost exclusively from Mexican American census tracts in San Antonio, Texas. The major objectives of this study are to identify low frequency or rare variants in and around known common variant signals for CVD, as well as to find novel low frequency or rare variants influencing susceptibility to CVD^28^. Whole genome sequencing (WGS) for the Trans-Omics in Precision Medicine (TOPMed) program was supported by the National Heart, Lung and Blood Institute (NHLBI). Collection of the San Antonio Family Study data was supported in part by National Institutes of Health (NIH) grants R01 HL045522, MH078143, MH078111 and MH083824; and whole genome sequencing of SAFS subjects was supported by U01 DK085524 and R01 HL113323. We are very grateful to the participants of the San Antonio Family Study for their continued involvement in our research programs.

**Samoan Adiposity Study (SAS, n=1,182):**

*TOPMed dbGaP accession#: phs000972, Parent dbGaP accession#: phs000914,*

*Sequencing Center: New York Genome Center and McDonnell Genome Institute (MGI) at Washington University.*

The main aim of the Samoan Adiposity Study is to identify risk factors for obesity and cardiometabolic phenotypes among the Samoan population. The recruitment and measurement of the study participants took place from February to July 2010 and thirty-three villages were included in the study^29,30^. The participants reside throughout the independent nation of Samoa, which is experiencing economic development and the nutrition transition. Genotyping was performed with the Affymetrix Genome-Wide Human SNP 6.0 Array using a panel of approximately 900,000 SNPs. Anthropometric, fasting blood biomarkers and detailed dietary, physical activity, health and socio-demographic variables were collected. Whole genome sequencing (WGS) for the Trans-Omics in Precision Medicine (TOPMed) program was supported by the National Heart, Lung and Blood Institute (NHLBI). Data collection was funded by NIH grant R01-HL093093. We thank the Samoan participants of the study and local village authorities. We acknowledge the support of the Samoan Ministry of Health and the Samoa Bureau of Statistics for their support of this research.

**Taiwan Study of Hypertension using Rare Variants (THRV, n=1,982):**

*TOPMed dbGaP accession#: phs001387, Parent dbGaP accession#: phs001387, Sequencing Center: Baylor College of Medicine Human Genome Sequencing Center.*

The THRV-TOPMed study consists of three cohorts: The SAPPHIRe Family cohort, TSGH (Tri-Service General Hospital, a hospital-based cohort), and TCVGH (Taichung Veterans General Hospital, another hospital-based cohort), all based in Taiwan^31,32^. 1,271 subjects were previously recruited as part of the NHLBI-sponsored SAPPHIRe Network (which is part of the Family Blood Pressure Program, FBPP). THRV is a collaborative study between Washington University in St. Louis, LA BioMed at Harbor UCLA, University of Texas in Houston, Taichung Veterans General Hospital, Taipei Veterans General Hospital, Tri-Service General Hospital, National Health Research Institutes, National Taiwan University, and Baylor University. THRV is based (substantially) on the parent SAPPHIRe study, along with additional population-based and hospital-based cohorts. Whole genome sequencing (WGS) for the Trans-Omics in Precision Medicine (TOPMed) program was supported by the National Heart, Lung and Blood Institute (NHLBI). The Rare Variants for Hypertension in Taiwan Chinese (THRV) is supported by the National Heart, Lung, and Blood Institute (NHLBI) grant (R01HL111249) and its participation in TOPMed is supported by an NHLBI supplement (R01HL111249-04S1). SAPPHIRe was supported by NHLBI grants (U01HL54527, U01HL54498) and Taiwan funds, and the other cohorts were supported by Taiwan funds.

**Women’s Health Initiative (WHI, n=8,263):**

*TOPMed dbGaP accession#: phs001237, Parent dbGaP accession#: phs000200, Sequencing Center: Broad Institute of MIT and Harvard.*

The Women's Health Initiative (WHI) is a large study of postmenopausal women’s health investigating risk factors for cancer, CVD, age-related fractures and chronic disease. It began in 1993 as a set of randomized controlled clinical trials (CT) and an observational study (OS). Specifically, the CT (n=68,132) included three overlapping components: The Hormone Therapy (HT) Trials (n=27,347), Dietary Modification (DM) Trial (n=48,835), and Calcium and Vitamin D (CaD) Trial (n=36,282). Eligible women could be randomized into as many as all three CTs components. Women who were ineligible or unwilling to join the CT were then invited to join the OS (n=93,676)^33,34^. WHI is a case control study with ~5000 strokes and >2000 CHD samples. Whole genome sequencing (WGS) for the Trans-Omics in Precision Medicine (TOPMed) program was supported by the National Heart, Lung and Blood Institute (NHLBI). The WHI program is funded by the National Heart, Lung, and Blood Institute, National Institutes of Health, U.S. Department of Health and Human Services through contracts HHSN268201600018C, HHSN268201600001C, HHSN268201600002C, HHSN268201600003C, and HHSN268201600004C.

**TOPMed Omics Support table:**

| **TOPMed Accession #** | **TOPMed Project** | **Parent Study** | **TOPMed Phase** | **Omics Center** | **Omics Support** | **Omics Type** |
| --- | --- | --- | --- | --- | --- | --- |
| **phs000956** | Amish | Amish | 1 | Broad Genomics | 3R01HL121007-01S1 | WGS |
| **phs001211** | AFGen | ARIC AFGen | 1 | Broad Genomics | 3R01HL092577-06S1 | WGS |
| **phs001211** | VTE | ARIC | 2 | Baylor | 3U54HG003273-12S2 / HHSN268201500015C | WGS |
| **phs001644** | AFGen | BioMe AFGen | 2.4 | MGI | 3UM1HG008853-01S2 | WGS |
| **phs001644** | BioMe | BioMe | 3 | MGI | HHSN268201600037I | WGS |
| **phs001644** | BioMe | BioMe | 3 | Baylor | HHSN268201600033I | WGS |
| **phs001612** | CARDIA | CARDIA | 3 | Baylor | HHSN268201600033I | WGS |
| **phs000954** | CFS | CFS | 3.5 | NWGC | HHSN268201600032I | WGS |
| **phs000954** | CFS | CFS | 1 | NWGC | 3R01HL098433-05S1 | WGS |
| **phs001368** | CHS | CHS | 3 | Baylor | HHSN268201600033I | WGS |
| **phs001368** | CHS | CHS | 5.5-5.6; 6 backfill | Broad Genomics | HHSN268201600034I | WGS |
| **phs001368** | VTE | CHS VTE | 2 | Baylor | 3U54HG003273-12S2 / HHSN268201500015C | WGS |
| **phs001412** | AA_CAC | DHS | 2 | Broad Genomics | HHSN268201500014C | WGS |
| **phs000974** | AFGen | FHS AFGen | 1 | Broad Genomics | 3R01HL092577-06S1 | WGS |
| **phs000974** | FHS | FHS | pilot; 4.5; 5.5-5.6; 6 backfill | Broad Genomics | HHSN268201600034I | WGS |
| **phs000974** | FHS | FHS | 1 | Broad Genomics | 3U54HG003067-12S2 | WGS |
| **phs001218** | AA_CAC | GeneSTAR AA_CAC | 2 | Broad Genomics | HHSN268201500014C | WGS |
| **phs001218** | GeneSTAR | GeneSTAR | 2 | Psomagen | 3R01HL112064-04S1 | WGS |
| **phs001218** | GeneSTAR | GeneSTAR | legacy | Illumina | R01HL112064 | WGS |
| **phs001345** | HyperGEN_GENOA | GENOA | 2 | NWGC | 3R01HL055673-18S1 | WGS |
| **phs001345** | AA_CAC | GENOA AA_CAC | 2 | Broad Genomics | HHSN268201500014C | WGS |
| **phs001217** | GenSalt | GenSalt | 2 | Baylor | HHSN268201500015C | WGS |
| **phs001359** | GOLDN | GOLDN | 2 | NWGC | 3R01HL104135-04S1 | WGS |
| **phs001395** | HCHS_SOL | HCHS_SOL | 3 | Baylor | HHSN268201600033I | WGS |
| **phs001293** | HyperGEN_GENOA | HyperGEN | 2 | NWGC | 3R01HL055673-18S1 | WGS |
| **phs000964** | JHS | JHS | 1 | NWGC | HHSN268201100037C | WGS |
| **phs001416** | AA_CAC | MESA AA_CAC | 2 | Broad Genomics | HHSN268201500014C | WGS |
| **phs001416** | MESA | MESA | 2 | Broad Genomics | 3U54HG003067-13S1 | WGS |
| **phs001062** | AFGen | MGH_AF | 1 | Broad Genomics | 3R01HL092577-06S1 | WGS |
| **phs001062** | AFGen | MGH_AF | 1.4; 1.5; 2.4 | Broad Genomics | 3U54HG003067-12S2 / 3U54HG003067-13S1; 3U54HG003067-12S2 / 3U54HG003067-13S1; 3UM1HG008895-01S2 | WGS |
| **phs001215** | SAFS | SAFS | legacy | Illumina | R01HL113322 | WGS |
| **phs001215** | SAFS | SAFS | 1 | Illumina | 3R01HL113323-03S1 | WGS |
| **phs000972** | Samoan | Samoan | 2 | NYGC Genomics | HHSN268201500016C | WGS |
| **phs000972** | Samoan | Samoan | 1 | NWGC | HHSN268201100037C | WGS |
| **phs001387** | THRV | THRV | 2 | Baylor | 3R01HL111249-04S1 / HHSN26820150015C | WGS |
| **phs001237** | WHI | WHI | 2 | Broad Genomics | HHSN268201500014C | WGS |

- Baylor = Baylor College of Medicine Human Genome Sequencing Center
- Broad Genomics = Broad Institute Genomics Platform
- Broad Metabolomics = Broad Institute and Beth Israel Metabolomics Platform
- Illumina = Illumina
- Keck MGC = Keck Molecular Genomics Core Facility
- MGI = McDonnell Genome Institute
- NWGC = Northwest Genomics Center
- NYGC Genomics = New York Genome Center Genomics
- Psomagen = Psomagen

**Supplementary Figure Legends:**

**Supplementary Fig. 1:**

**Overview of the workflow implement in the study.** Multi-ancestral whole genome sequenced data from TOPMed freeze 8, including 66329 samples were qualified for the analysis from five different ancestral groups, were analyzed with lipids. Single variant GWAS and genome-wide rare variant aggregates tests were conducted using SAIGE and STAAR methods, respectively. The results were compared with the GWAS catalog and MVP summary stats to identify novel associations. Conditional analyses of known associated single variants were conducted for rare variant association tests at known loci. Mendelian lipid loci were explored in further detail.

TOPMed – Trans-Omics for Precision Medicine; GWAS – Genome Wide Association Study; SAIGE – Scalable and Accurate Implementation of GEneralized mixed model; STAAR – variant-Set Test for Association using Annotation information; MVP – Million Veteran Program

**Supplementary Fig. 2:**

**Manhattan plots for the four lipid phenotypes.** MH plots for single variant GWAS for the four lipids. Genes nearby to the variants that are strongly associated at a locus are mapped on the plot for each chromosome. Genes linked to SNPs at potentially novel loci in TOPMed are noted in red. **a**, MH plots for HDL-C association. **b**, MH plots for LDL-C association. **c**, MH plots for TC association. **d**, MH plots for TG association.

MH – Manhattan; GWAS – Genome Wide Association Study; SNPs – Single Nucleotide Polymorphisms; TOPMed – Trans-Omics for Precision Medicine; HDL-C – High-Density Lipoprotein Cholesterol; LDL-C – Low-Density Lipoprotein Cholesterol; TC – Total Cholesterol; TG – Triglycerides.

**Supplementary Fig. 3:**

**Correlation of *CETP* gene expression effects with HDL-C effects**. The effects from GWAS and GTEx data from Liver, Adipose Subcutaneous and Adipose Visceral tissues for the eQTLs were correlated. The x-axis contains each gene linked to the eQTLs and y-axis represents the correlation coefficients, where the x-axis is ordered based on the correlation coefficients and size of each datapoint is based on the p-value of the correlation test. *CETP* along with other genes have a strong negative correlation of the effect estimates. **a**, Liver tissue data **b**, Adipose Subcutaneous tissue data **c**, Adipose Visceral (Omentum) tissue data.

GWAS – Genome Wide Association Study; GTEx – Genotype-Tissue Expression; eQTLs – Expression Quantitative Trait Loci; HDL-C – High-Density Lipoprotein Cholesterol.

**Supplementary Fig. 4**

**Genome wide view of coding and non-coding aggregate sets.** Aggregate sets that were significant with STAAR-O p-values lesser than 5E-04 were retained. The genes are ordered based on chromosomes, following by their genomic positions. The coloring scale is based on -log10(STAAR-O p-values). The most significant mask has bright red tile. Aggerate sets significant after conditional analysis is represented with the corresponding genes on the right panel, for coding and non-coding aggregates. **a**, Rare variant aggregates for HDL-C. **b**, Rare variant aggregates for LDL-C. **c**, Rare variant aggregates for TC. **d**, Rare variant aggregates for TG.

STAAR – variant-Set Test for Association using Annotation information; HDL-C – High-Density Lipoprotein Cholesterol; LDL-C – Low-Density Lipoprotein Cholesterol; TC – Total Cholesterol; TG – Triglycerides.

**Supplementary Fig. 5**

**Conditional analysis of coding rare-variants from the same gene and a near-by gene**. Non-coding rare variant sets significantly associated with HDL-C and LDL-C after the conditional analysis on known variants are shown with additional adjustment on rare-coding variants. The different colored dots on the plot represents the conditional STAAR-O p-values when adjusting for known variants (Set1) and rare-coding variants of the same or near-by gene. **a**, Non-coding rare variant sets significantly associated with HDL-C. Conditional analysis shows that the signal from *APOC3* and *APOA1* drop after it was adjusted for coding rare-variants. Whereas the signal from *CETP* locus combined with *HERPUD1* variant sets are retained. Similarly, *GFOD2* non-coding rare-variant aggregate is significant, where there is no change in STAAR-O p-values after the conditional analysis. The dotted line separates the variants sets that are significant and insignificant after the additional conditional analysis. **b**, Non-coding rare variant sets significantly associated with LDL-C. After adjustment, *APOB* signal drops minimally, whereas *PCSK9* signal enhances. Conditioning on the near-by gene *LDLR*, the signal from *SPC24* enhancer-DHS improves, but enhancer-CAGE drops significantly.

STAAR – variant-Set Test for Association using Annotation information; HDL-C – High-Density Lipoprotein Cholesterol; LDL-C – Low-Density Lipoprotein Cholesterol; CAGE – Cap Analysis of Gene Expression; DHS – DNase hypersensitivity.

**Supplementary Fig. 6**

**Distribution of Mendelian gene STAAR-O p-values.** The 22 genes are ordered based on the lipid categories (HDL-C, LDL-C and TG) and the most significant association of each gene aggregate set is represented as p-values. **a**, -log10(STAAR-O p-values) of the five different coding masks. **b**, -log10(STAAR-O p-values) of the seven different non-coding masks.

STAAR – variant-Set Test for Association using Annotation information; LDL-C – Low-Density Lipoprotein Cholesterol; TG – Triglycerides.

**Supplementary Table Legends:**

**Supplementary Table 1:**

**TOPMed freeze 8 phenotype data distributed based on 21 cohorts**. The Freeze8 lipid data is composed of 66329 samples. Samples are tabulated based on gender, ancestral groups, sequencing centers and lipid treatment (percentage of samples) for each cohort. Distribution of mean (standard deviation) for sex, unadjusted non-transformed lipid concentration of HDL-C, LDL-C, TC and TG in full sample group and stratified groups based on gender and ancestry are provided. Since TG has a skewed distribution, the median and IQR for TG concentration are provided separately for all samples. Lipid concentrations are in units of mg/dl. Cohorts with no values for any specific columns are represented as NA. Sequencing centers: Baylor: Baylor College of Medicine Human Genome Sequencing Center, Broad: Broad Institute of MIT and Harvard, Illumina: Illumina Genomic Services, Macrogen: PSOMAGEN (formerly Macrogen), NYGC: New York Genome Center, UW: McDonnell Genome Institute (MGI) at Washington University, WASHU: Northwest Genomics Center.

HDL-C – High-Density Lipoprotein Cholesterol; LDL-C – Low-Density Lipoprotein Cholesterol; TC – Total Cholesterol; TG – Triglycerides

**Supplementary Table 2:**

**TOPMed Freeze 8 genotype distribution based on chromosome.** Variant counts by chromosome are tabulated across MAF and MAC bins.

MAF – Minor Allele Frequency; MAC – Minor Allele Count

**Supplementary Table 3:**

**TOPMed Freeze 8 variants summary.** Variants which passed the significance criteria were clumped (window 250 kb, r^2^ 0.5) and compared against MVP summary statistic and GWAS catalog. Variants were binned to three categories, Known-Position (variant previously associated), Known-Loci (variants not previously significantly associated with the corresponding lipid phenotype but within 500 kb of a known locus) and Novel. The list of variants is tabulated for each lipid phenotype and each category of is ordered based on chromosome position.

TOPMed – Trans-Omics for Precision Medicine; MVP – Million Veteran Program; GWAS – Genome Wide Association Study.

**Supplementary Table 4:**

**Ancestry specificity and replication of putative novel variants.** List of novel single variants identified after comparing with MVP summary stats are tabulated. Effects, p-values MAF of variants in discovery (TOPMed) cohort specific to each ancestral group is documented**.** Effects and p-values from replication cohort (MGB and Penn biobank) are documented. All effect estimates are in mg/dL units, except for TG which was log-transformed in analysis thereby representing fractional change.

MVP – Million Veteran Program; MAF – Minor Allele Frequency; TOPMed – Trans-Omics for Precision Medicine; MGB – Mass General Brigham.

**Supplementary Table 5:**

**Baseline characteristics of replication cohorts.** Sample sizes, gender distributions, ancestry distributions, and mean ages of individuals in each of the replication cohort are provided. Mean (standard deviation) of unadjusted and nontransformed lipid concentrations are presented for the MGB Biobank and Penn Medicine Biobank.

MGB – Mass General Brigham.

**Supplementary Table 6:**

**Evaluation of suggestive lipid alleles in TOPMed with independent datasets.** Putative novel variants at ‘suggestive’ p-values (5x10^-07^ – 5x10^-09^) in TOPMed are listed, including ancestry-specific effects. Evidence for association in the MGB and Penn Medicine Biobanks are also included.

TOPMed – Trans-Omics for Precision Medicine; MGB – Mass General Brigham.

**Supplementary Table 7:**

**List of *CETP* variants significant at suggestive p-value (5X10^-07^) from both LDL-C and HDL-C GWAS**. All the variants were highly significant to HDL-C and the White ancestry group contributes mostly for the association with positive effects. Three variants are genome significant (5X10^-09^) and two of them are suggestive significant with LDL-C association. The Black ancestry group contributed to strong LDL-C association with negative effects. Effects, p-values and MAF of trans-ancestry and ancestry-specific associations are documented.

HDL-C – High-Density Lipoprotein Cholesterol; LDL-C – Low-Density Lipoprotein Cholesterol; GWAS – Genome Wide Association Studies; MAF – Minor Allele Frequency.

**Supplementary Table 8:**

**Significant gene-centric coding rare variant aggregate sets.** Each of the significant (p-value < 2.5x10^-06^) rare (MAF<1%) coding aggregate sets for at least one lipid phenotype are listed by gene name and mask. Aggregates are ordered by mask and number of variants in each set is documented. SKAT, Burden, ACAT-O and STAAR-O p-values for each of the aggregate sets are provided.

MAF – Minor Allele Frequency; SKAT – SNP-set (Sequence) Kernel Association Test; ACAT – Aggregated Cauchy Association Test; STAAR – variant-Set Test for Association using Annotation information.

**Supplementary Table 9:**

**Significant gene-centric non-coding rare variant aggregate sets.** Significant (p-value < 2.5x10^-06^) rare (MAF<1%) non-coding aggregate sets for at least one lipid phenotype are listed. Aggregates are ordered by masks and the number of variants in each aggregate set is documented. SKAT, Burden, ACAT-O and STAAR-O p-values for each of the aggregate sets are provided.

MAF – Minor Allele Frequency; SKAT – SNP-set (Sequence) Kernel Association Test; ACAT – Aggregated Cauchy Association Test; STAAR – variant-Set Test for Association using Annotation information.

**Supplementary Table 10:**

**Significant gene-centric coding rare variant aggregate sets after conditional analysis.** Each of the rare coding aggregate sets significant genome wide (after conditional analysis on known common variants) are listed for each lipid phenotypes. The list of common variants that were used for adjustment is provided with RS ids. Aggregates are ordered based on STAAR-O p-values and the number of variants in each aggregate set is documented. SKAT, Burden, ACAT-O and STAAR-O p-values for each of the aggregate sets are provided. The effect estimates and corresponding p-values from Glmm.Wald test is documented.

SKAT – SNP-set (Sequence) Kernel Association Test; ACAT – Aggregated Cauchy Association Test; STAAR – variant-Set Test for Association using Annotation information.

**Supplementary Table 11:**

**Significant gene-centric non-coding rare variant aggregate sets after conditional analysis.** Each of the rare non-coding aggregate sets significant genome wide (after conditional analysis on known common variants) are listed for each lipid phenotypes. The list of common variants that were used for adjustment is provided with RS ids. Aggregates are ordered based on STAAR-O p-values and the number of variants in each aggregate set is documented. SKAT, Burden, ACAT-O and STAAR-O p-values for each of the aggregate sets are provided. The effect estimates and corresponding p-values from Glmm.Wald test is documented.

SKAT – SNP-set (Sequence) Kernel Association Test; ACAT – Aggregated Cauchy Association Test; STAAR – variant-Set Test for Association using Annotation information.

**Supplementary Table 12:**

**Region-based sliding-window results before and after conditional analysis.** Regions that are significantly associated with at least one lipid phenotype after conditioning on significant single variants are listed. The start and end location of the region, the number of variants comprising the aggregate sets and the list of variants which were conditioned are provided. Aggregates are ordered based on STAAR-O p-values and the number of variants in each aggregate set is documented. SKAT, Burden, ACAT and STAAR-O p-values before and after conditional analysis for each of the aggregate sets are provided. Genes mapped to the region and the intron-exon boundaries are provided. Glmm.Wald test was implemented on all the variants of the aggregate set and only non-coding variants for each aggregate. The corresponding effect estimates, and p-values is documented.

SKAT – SNP-set (Sequence) Kernel Association Test; ACAT – Aggregated Cauchy Association Test; STAAR – variant-Set Test for Association using Annotation information.

**Supplementary Table 13:**

**Region-based dynamic-window results before and after conditional analysis.** Regions that are significantly associated with at least one lipid phenotype by SCANG after conditioning on significant single variants are listed. The start and end location of the region, the number of variants comprising the aggregate sets and the list of variants which were conditioned are provided. Aggregates are ordered based on STAAR-O p-values and the number of variants in each aggregate set is documented. SKAT, Burden, ACAT and STAAR-O p-values before and after conditional analysis for each of the aggregate sets are provided. Unconditional p-values are obtained from each of the test where the aggregate set is significant, and the conditional p-values are summarized for each aggregate set. Genes mapped to the regions and the intron-exon boundaries are provided. Glmm.Wald test was implemented on all the variants of the aggregate set and only non-coding variants for each aggregate. The corresponding effect estimates, and p-values is documented.

SKAT – SNP-set (Sequence) Kernel Association Test; ACAT – Aggregated Cauchy Association Test; STAAR – variant-Set Test for Association using Annotation information.

**Supplementary Table 14:**

**Significant rare non-coding variants after conditioning on rare coding variants.** Rare non-coding aggregates sets were additionally adjusted for rare coding variants of the same gene and the nearby gene. The nearby gene pairs are as follows: *LDLR-SPC24*, *CETP-HERPUD1*, *APOC3-APOA1*. STAAR-O p-values at each adjustment step is provided and aggregates tested with nearby gene is additionally shown.

**Supplementary Table 15:**

**Common variants and rare variant aggregates at Mendelian lipid genes.** Twenty-two Mendelian lipid genes and their common variant and rare variant aggregates from coding and non-coding tests are listed.

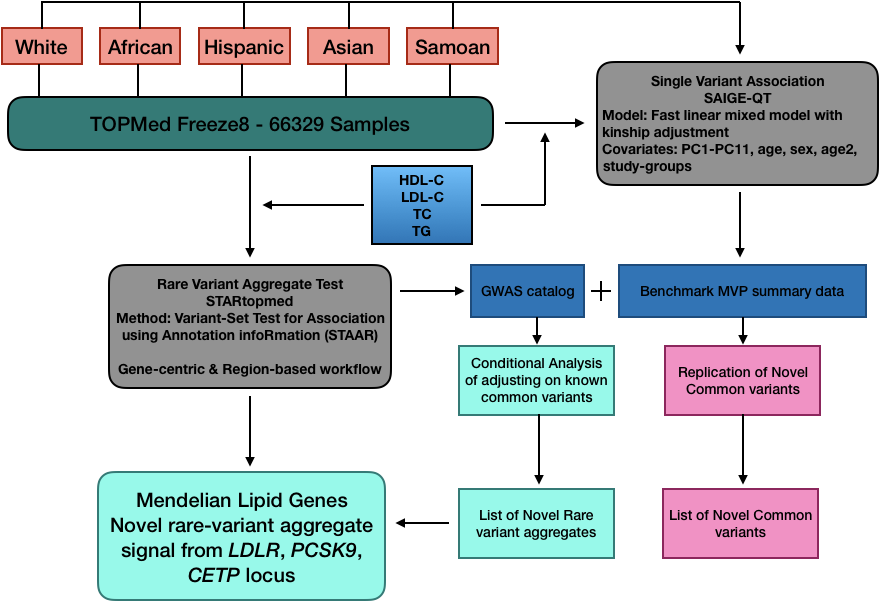

**Supplementary Fig. 1**

**Overview of the workflow implement in the study.**

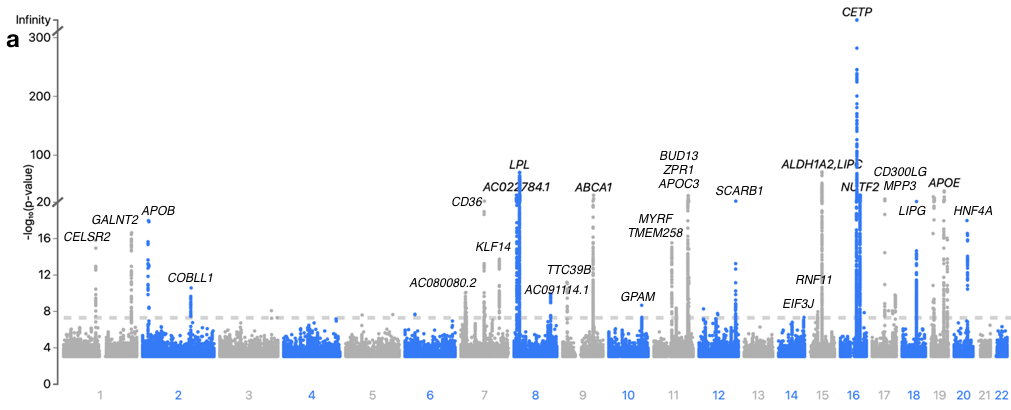

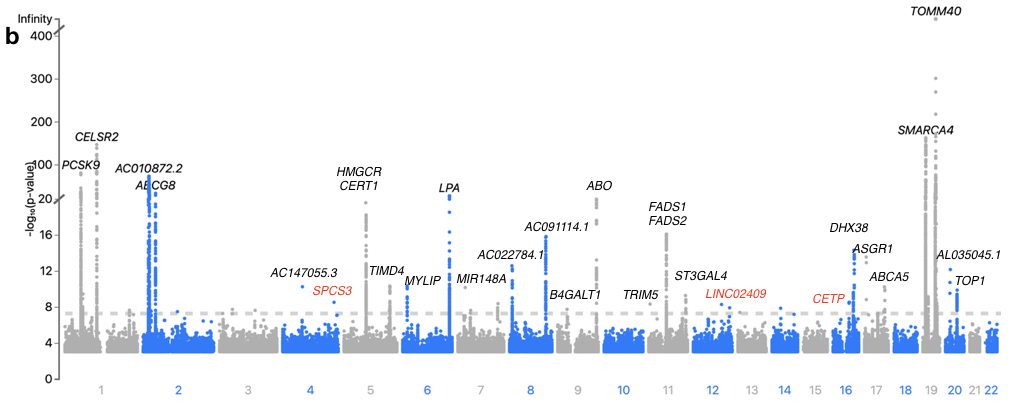

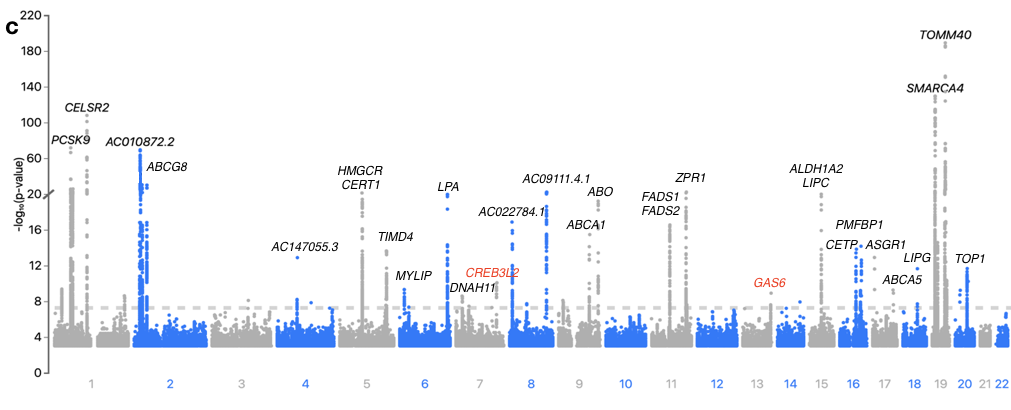

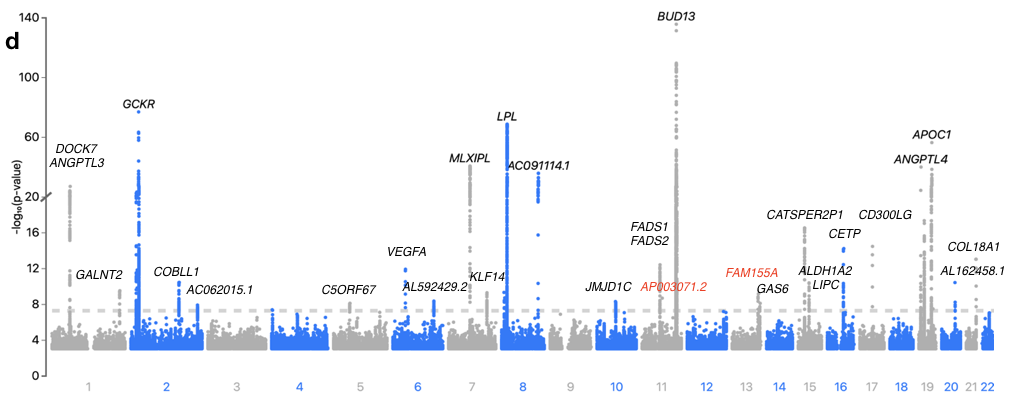

**Supplementary Fig. 2**

**Manhattan plots for the four lipid phenotypes.**

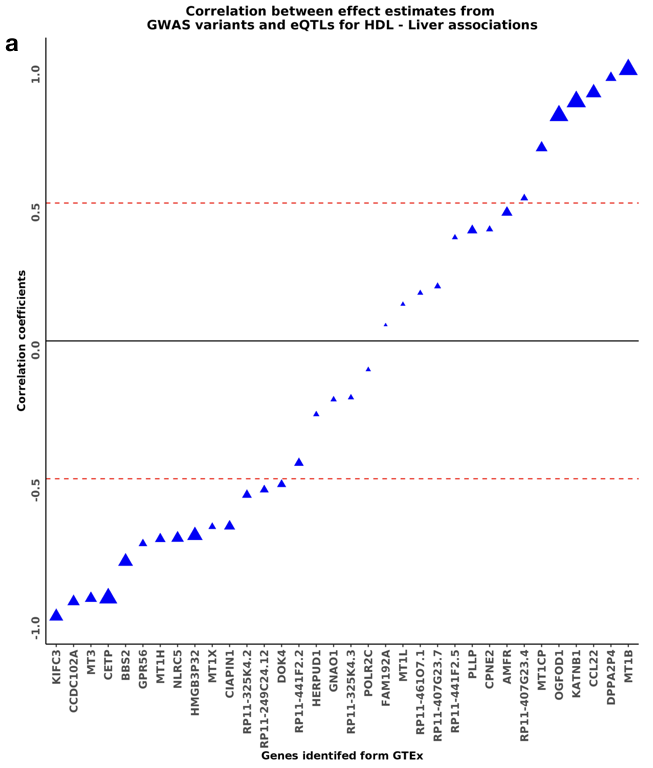

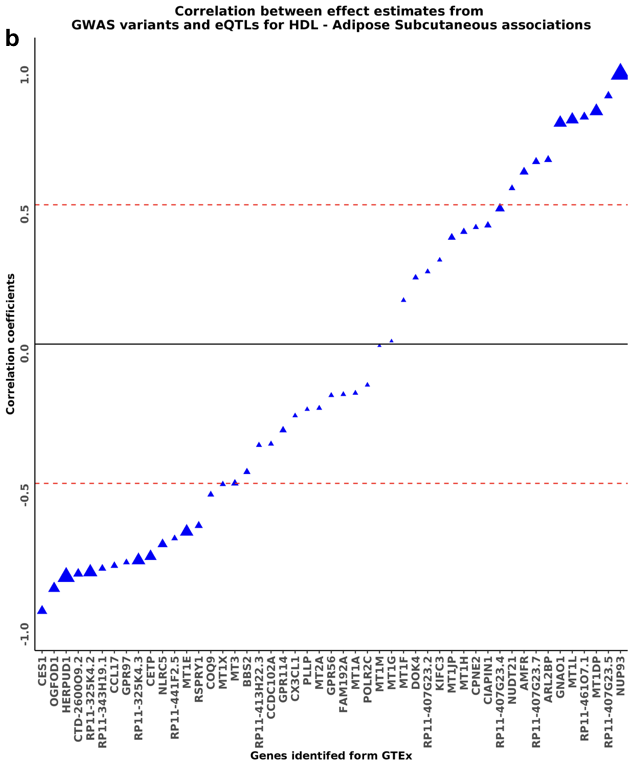

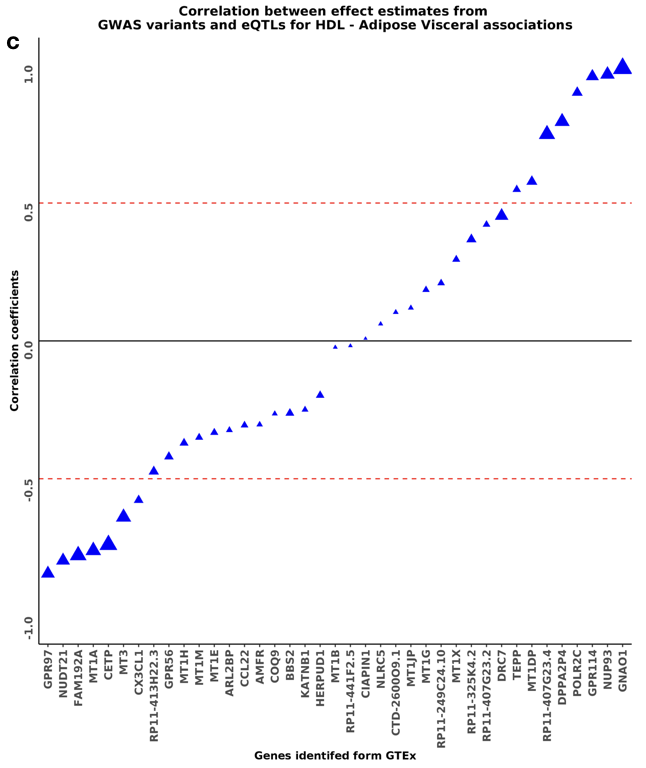

**Supplementary Fig. 3**

**Correlation of *CETP* gene expression effects with HDL-C effects**.

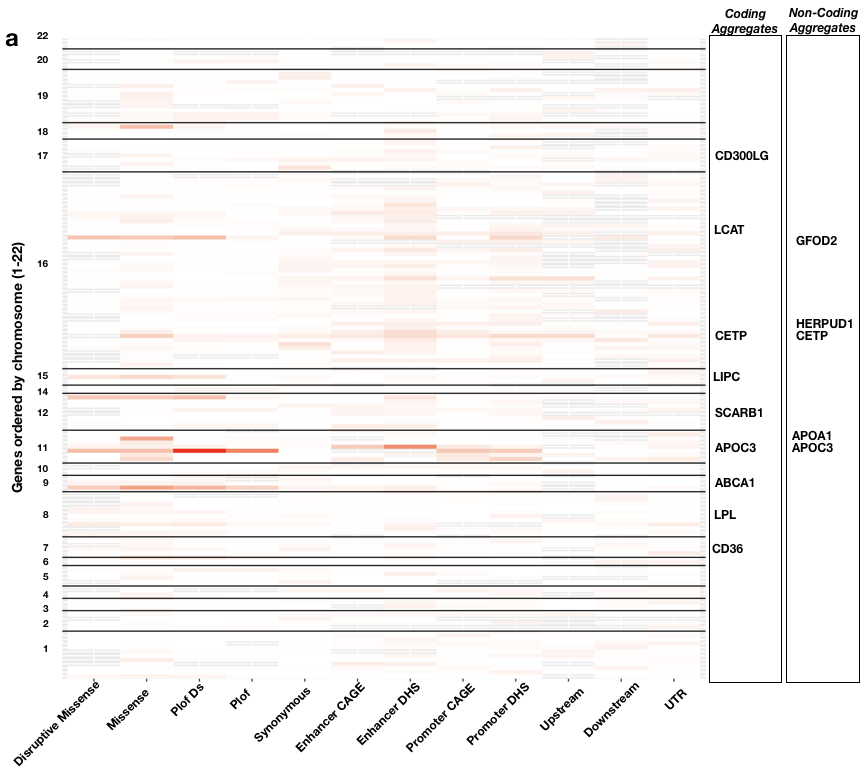

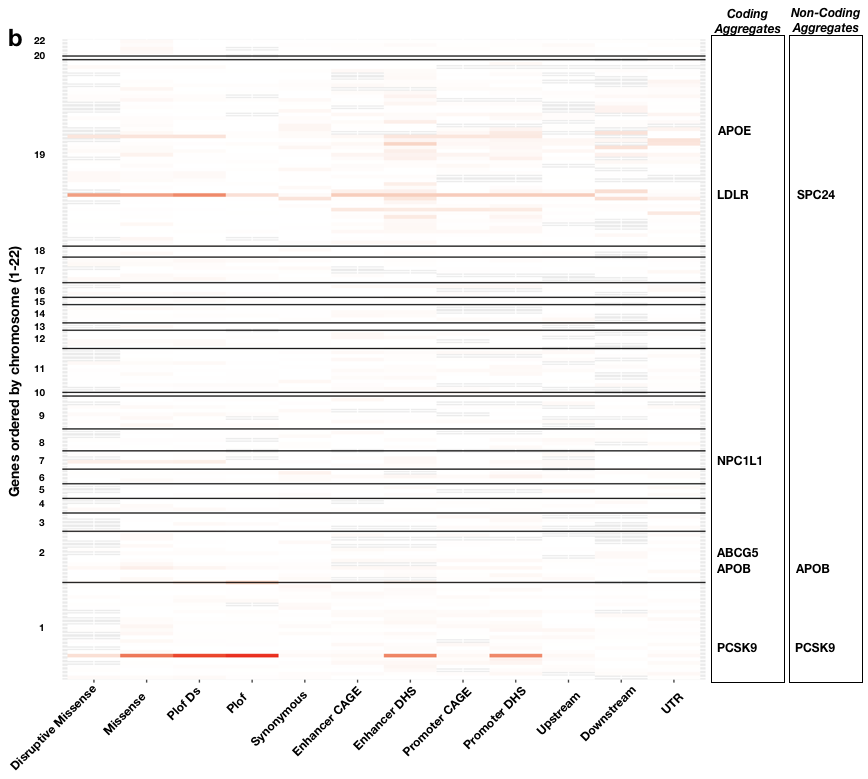

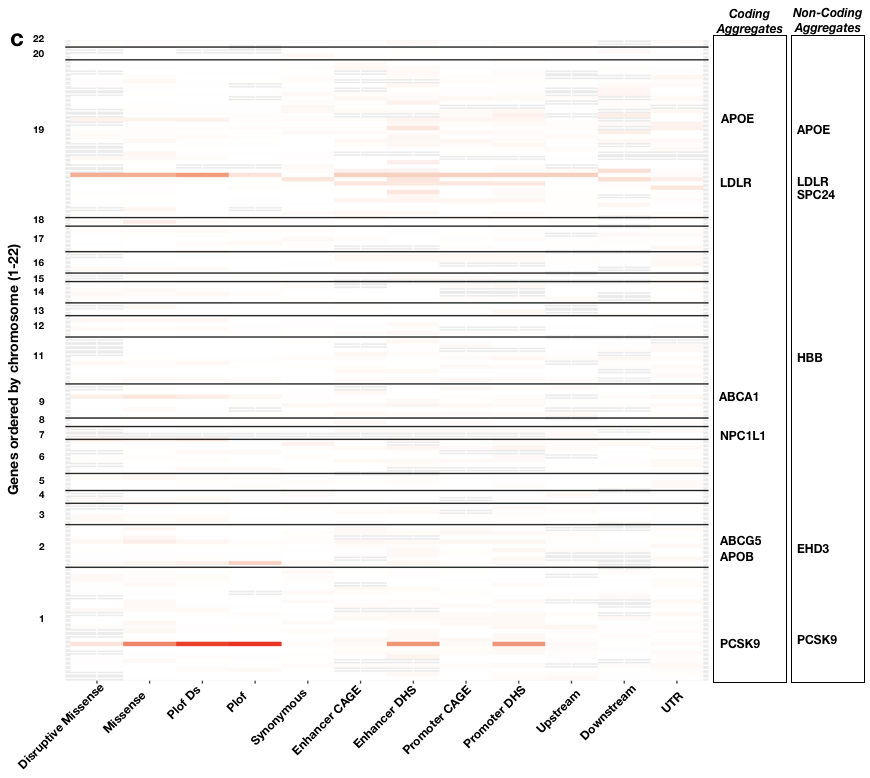

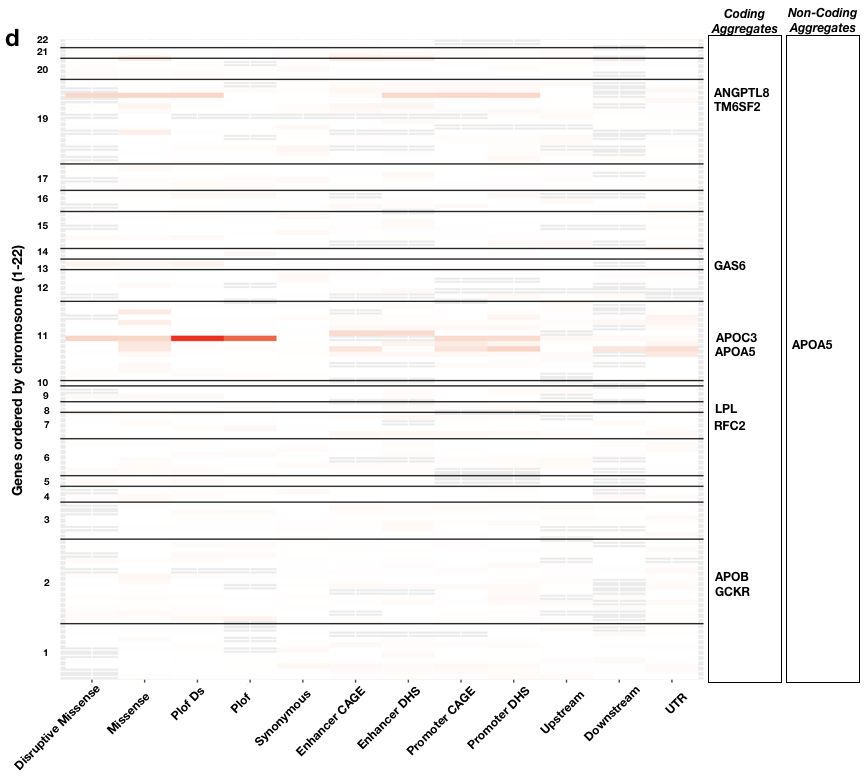

**Supplementary Fig. 4**

**Genome wide view of coding and non-coding aggregate sets.**

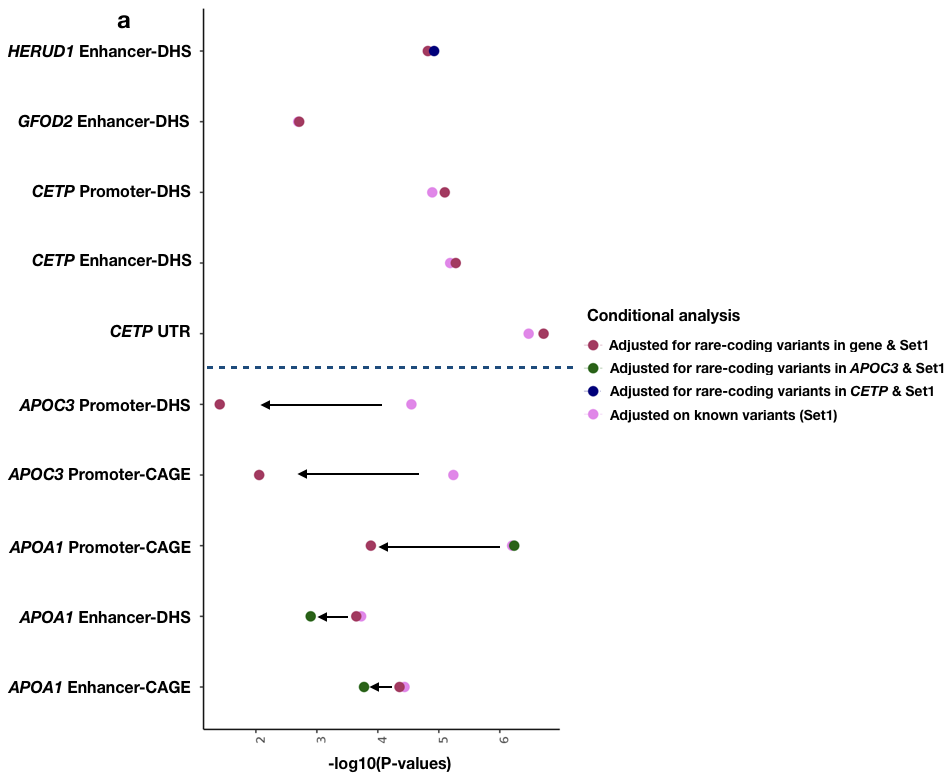

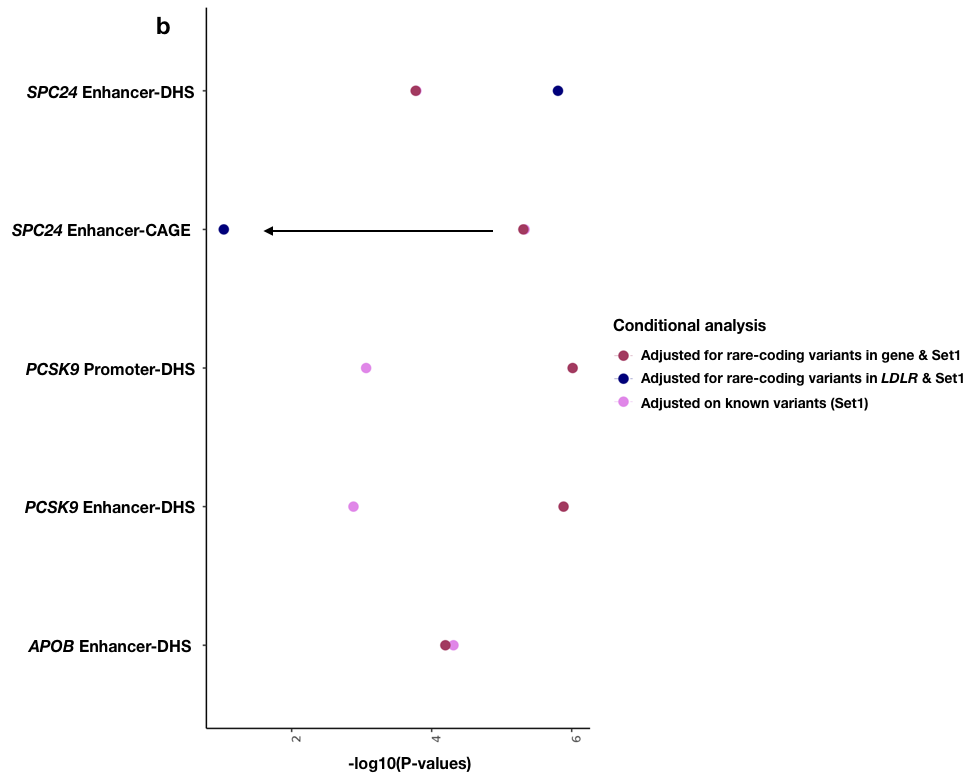

**Supplementary Fig. 5**

**Conditional analysis of coding rare-variants from the same gene and a near-by gene**.

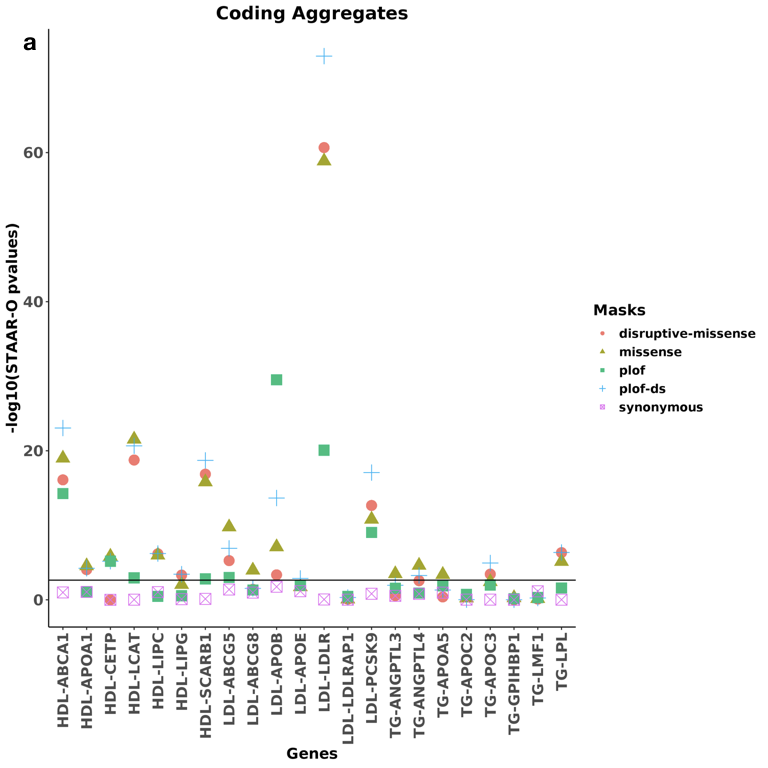

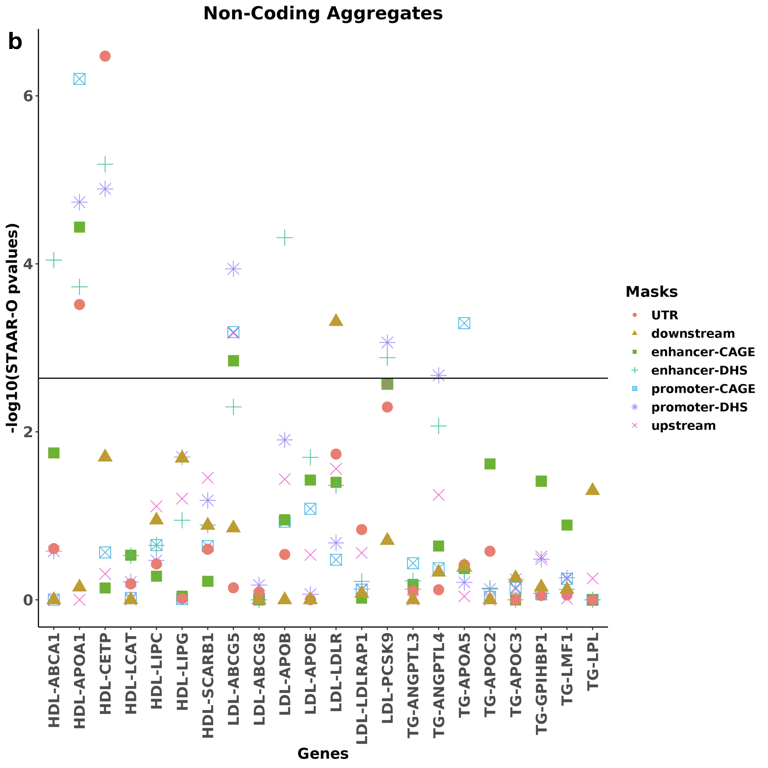

**Supplementary Fig. 6**

**Distribution of Mendelian gene STAAR-O p-values.**
